# A consensus phylogenomic approach highlights paleopolyploid and rapid radiation in the history of Ericales

**DOI:** 10.1101/816967

**Authors:** Drew A. Larson, Joseph F. Walker, Oscar M. Vargas, Stephen A. Smith

## Abstract

**Premise of study:** Large genomic datasets offer the promise of resolving historically recalcitrant species relationships. However, different methodologies can yield conflicting results, especially when clades have experienced ancient, rapid diversification. Here, we analyzed the ancient radiation of Ericales and explored sources of uncertainty related to species tree inference, conflicting gene tree signal, and the inferred placement of gene and genome duplications.

**Methods:** We used a hierarchical clustering approach, with tree-based homology and orthology detection, to generate six filtered phylogenomic matrices consisting of data from 97 transcriptomes and genomes. Support for species relationships was inferred from multiple lines of evidence including shared gene duplications, gene tree conflict, gene-wise edge-based analyses, concatenation, and coalescent-based methods and is summarized in a consensus framework.

**Key Results:** Our consensus approach supported a topology largely concordant with previous studies, but suggests that the data are not capable of resolving several ancient relationships due to lack of informative characters, sensitivity to methodology, and extensive gene tree conflict correlated with paleopolyploidy. We found evidence of a whole genome duplication before the radiation of all or most ericalean families and demonstrate that tree topology and heterogeneous evolutionary rates impact the inferred placement of genome duplications.

**Conclusions:** Our approach provides a novel hypothesis regarding the history of Ericales and confidently resolves most nodes. We demonstrate that a series of ancient divergences are unresolvable with these data. Whether paleopolyploidy is a major source of the observed phylogenetic conflict warrants further investigation.

## INTRODUCTION

The flowering plant clade Ericales contains several ecologically important lineages that shape the structure and function of ecosystems including tropical rainforests (e.g. Lecythidaceae, Sapotaceae, Ebenaceae), heathlands (e.g. Ericaceae), and open habitats (e.g. Primulaceae, Polemoniaceae) around the globe (ter Steege et al., 2006; Hedwall et al., 2013; He et al., 2014; Memiaghe et al., 2016; Moquet et al., 2017). With 22 families comprising ca. 12,000 species (Chase et al., 2016; Stevens, 2001 onward), Ericales are a diverse and disparate clade with an array of economically and culturally important plants. These include agricultural crops such as blueberries (Ericaceae), kiwifruits (Actinidiaceae), sapotas (Sapotaceae), Brazil nuts (Lecythidaceae), and tea (Theaceae) as well as ornamental plants such as cyclamens and primroses (Primulaceae), rhododendrons (Ericaceae), and phloxes (Polemoniaceae). Parasitism has arisen multiple times in Ericales (Monotropoidiae:Ericaceae, Mitrastemonaceae), as has carnivory in the American pitcher plants (Sarraceniaceae). Although Ericales have been a well-recognized clade throughout the literature (Chase et al., 1993; Anderberg et al., 2002; Schönenberger et al., 2005; Rose et al., 2018, Leebens-Mack et al. 2019), the evolutionary relationships among major clades within Ericales remain contentious.

One of the first molecular studies investigating these deep relationships used three plastid and two mitochondrial loci; the authors concluded that the dataset was unable to resolve several interfamilial relationships (Anderber et al., 2002). These relationships were revisited by Schönenberger et al. (2005) with 11 loci (two nuclear, two mitochondrial, and seven chloroplast), who found that maximum parsimony and Bayesian analyses provided support for the resolution of some early diverging lineages. Rose et al. (2018) utilized three nuclear, nine mitochondrial, and 13 chloroplast loci in a concatenated supermatrix consisting of 49,435 aligned sites and including 4,531 ericalean species but with 87.6% missing data. Despite the extensive taxon sampling utilized in Rose et al. (2018), several relationships were only poorly supported, including several deep divergences that the authors show to be the result of an ancient, rapid radiation. The 1KP initiative (Leebens-Mack et al. 2019) analyzed trascriptomes from across green plants, including 25 species of Ericales; their results suggest that whole genome duplications (WGDs) have occurred several times within Ericales, but the equivocal support they recover at several deep ericalean nodes highlights the need for a more throughout investigation for conflicting interfamilial relationships within the group and the biological and methodological sources of phylogenetic incongruence (Anderber et al., 2002; Bremer et al., 2002; Schönenberger et al., 2005; Rose et al., 2018; Leebens-Mack et al., 2019).

Despite the increasing availability of genome-scale datasets, many relationships across the Tree of Life remain controversial, as research groups recover different answers to the same evolutionary questions, often with seemingly strong support (e.g. Shen et al., 2017). One benefit of genome-scale data for phylogenetics (i.e. phylogenomics) is the ability to examine conflicting signal within and among datasets, which can be used to help understand conflicting species tree results and conduct increasingly comprehensive investigations as to why some relationships remain elusive. A key finding in the phylogenomics literature has been the high prevalence of conflicting phylogenetic signal among genes at contentious nodes (e.g. Brown et al. 2017b; Reddy et al., 2017; Vargas et al., 2017; Walker et al., 2018a). Such conflict may be the result of biological processes (e.g., introgression, incomplete lineage sorting, horizontal gene transfer), but can also occur due to lack of phylogenetic information or other methodological artifacts (Richards et al., 2018; Gonçalves et al., 2019; Walker et al., 2019). By identifying regions of high conflict, it becomes possible to determine areas of the phylogeny where additional analyses are warranted and future sampling efforts might prove useful. Transcriptomes provide information from hundreds to thousands of coding sequences per sample and have elucidated many of the most contentious relationships in the green plant phylogeny (e.g. Simon et al., 2012; Wickett et al., 2014; Walker et al., 2017; Leebens-Mack et al., 2019). Transcriptomes also provide information about gene and genome duplications not provided by most other common sequencing protocols for non-model organisms. Gene duplications are often associated with important molecular evolutionary events in taxa, but duplicated genes are also inherited through descent and should therefore contain evidence for how clades are related. By leveraging the multiple lines of phylogenetic evidence and the large amount of data available from transcriptomes, several phylogenetic hypotheses can be generated and tested to gain a holistic understanding of contentious nodes in the Tree of Life.

In this study, we sought to understand the evolutionary history of Ericales by analyzing coding sequences from thousands of homolog clusters to investigate support for contentious interfamilial relationships, ancient gene and genome duplications, heterogeneous rates of evolution, and conflicting signal among genes. We examined the deep relationships of Ericales to determine whether the data strongly support any resolution. We include the possibility of there not being enough data to resolve these relationships (e.g., a hard or soft polytomy) despite previous resolutions and thousands of transcriptomic sequences. While future developments in methods and sampling will likely continue to elucidate many contentious relationships across the plant phylogeny, we consider whether a polytomy may represent a more justifiable representation of the evolutionary history of a clade than any single fully bifurcating species tree given our current resources.

In applying a phylogenetic consensus approach that considers several methodological alternatives we examined disagreement among methods. We explore this approach as a means of providing valuable information about whether the available data may be insufficient to confidently resolve a single, bifurcating species tree, even if a given methodology may suggest a resolved topology with strong support. Our investigation of the evolution of Ericales may be particularly well-suited to initiate a discussion about the impact of topological uncertainty on our ability to confidently resolve the placement of rare evolutionary events (e.g. whole genome duplication, major morphological innovation), the prevalence of biological polytomies across the Tree of Life, and when polytomies may be considered useful representations of evolutionary relationships in the postgenomic era.

## MATERIALS AND METHODS

### Taxon sampling and de novo assembly procedures

We assembled a dataset consisting of coding sequence data from 97 transcriptomes and genomes, the most extensive exome dataset for Ericales to date (Appendix S1; see Supplemental Data with this article). Samples assembled from raw reads were done so with Trinity version 2.5.1 using default parameters and the “--trimmomatic” option (Kodama et al., 2011; Grabherr et al., 2011; Matasci et al., 2014). Assembled reads were translated using TransDecoder v5.0.2 (Haas et al., 2013) with the option to retain BLAST hits to a custom protein database consisting of *Beta vulgaris*, *Arabidopsis thaliana*, and *Daucus carota* obtained from Ensembl plants (https://plants.ensembl.org/). The translated amino acid sequences from each assembly were reduced by clustering with CD-HIT version 4.6 with settings -c 0.995 -n 5 (Fu et al., 2012). As a quality control step, nucleotide sequences for the chloroplast genes *rbcL* and *matK* were extracted from each assembly using a custom script (see Data Accessibility Statement) and then queried using BLAST against the NCBI online database (Altschul et al., 1990). In cases where the top hits were to a species not closely related to that of the query, additional sequences were investigated and if contamination, misidentification, or other issues seemed likely, the transcriptome was not included in further analyses. The final transcriptome sampling included 86 ingroup taxa spanning 17 of the 22 families within Ericales as recognized in APG VI (Appendix S1; Chase et al., 2016).

### Homology inference

We used the hierarchical clustering procedure from Walker et al. (2018b) for homology identification. In short, the method involves performing an all-by-all BLAST procedure on user-defined clades; homolog clusters identified within each clade are then combined recursively with clusters of a sister clade based on sequence similarity until clusters from all clades have been combined. To assign taxa to groups for clustering, we identified coding sequences from the genes *rpoC2*, *rbcL*, *nhdF*, and *matK* using the same script as for the quality control step. Sequences from each gene were then aligned with MAFFT v7.271 (Katoh et al., 2002; Katoh and Standley, 2013). The alignments were concatenated using the command pxcat in phyx (Brown et al., 2017a) and the supermatrix was used to estimate a species tree with RAxML v8.2.12 (Stamatakis, 2014). The resulting tree was used to manually assign taxa to one of eight clades for initial homolog clustering (Appendix S2). Homolog clustering within each group was performed following the methods of Yang and Smith (2014).

Nucleotide sequences for the inferred homologs within each group were aligned with MAFFT v7.271 and columns with less than 10% occupancy were removed with the pxclsq command in phyx. Homolog trees were estimated with RAxML, unless the homolog had >500 tips, in which case FastTree v2.1.8 was used (Price et al., 2010). Tips with branch lengths longer than 1.5 substitutions/site were trimmed because the presence of highly divergent sequences in homolog clusters is often the result of misidentified homology (Yang and Smith, 2014). When a clade is formed of sequences from a single taxon, it likely represents either in-paralogs or alternative splice sites and since neither of these provide phylogenetic information, we retained only the tip with the longest sequence, excluding gaps introduced by alignment. This procedure was repeated with refined clusters twice using the same settings. Homologs among each of the eight groups were then recursively combined (https://github.com/jfwalker/Clustering) and homolog trees were again estimated and refined with the same settings except that internal branches longer than 1.5 substitutions/site were also cut—again to reduce potentially misidentified homology. Homolog trees were again re-estimated and refined by cutting terminal branches longer than 0.8 substitutions/site and internal branches longer than 1.0 substitutions/site. The final homolog set contained 9469 clusters.

### Initial ortholog identification and species tree estimation

Orthologs were extracted from homolog trees using the rooted tree (RT) method, which allows for robust orthology detection even after genome duplications (Yang and Smith, 2014). For the RT procedure, seven cornalean taxa as well as *Arabidopsis thaliana* and *Beta vulgaris* were used as outgroups (Appendix S1). Previous work has suggested that Ericales is sister to the euasterids with Cornales sister to Ericales+euasterids (Stevens, 2001 onwards), therefore we treated *Helianthus annuus* and *Solanum lycopersicum* as ingroup taxa for the purposes of the RT procedure so that orthologs were able to be rooted on a non-ericalean taxon after ortholog identification. Orthologs were not extracted from homolog groups with more than 5000 tips because of the uncertainty in reconstructing very large homolog trees (Walker et al., 2018b). Orthologs with sequences from fewer than 50 ingroup taxa were discarded to reduce the amount of missing data in downstream analyses. Because our interest is in addressing phylogenetic questions, rather than those of gene functionality, here we use the term ortholog to describe clusters of sequences that have been inferred to be monophyletic based on their position within inferred homolog trees after accounting for gene duplication. The tree-aware ortholog identification employed here should provide the best available safeguard against misidentified orthology that could mislead phylogenetic analyses (Eisen, 1998; Gabaldón, 2008; Yang and Smith, 2014; Brown and Thomson, 2016).

The resulting ortholog trees were then filtered to require at least one euasterid taxon, *Helianthus annuus* or *Solanum lycopersicum*, for use as an outgroup for rooting within each ortholog. If both outgroups were present in an ortholog tree but no bipartition existed with only those taxa (i.e. the tree could not be rooted on both), the ortholog was discarded because the monophyly of the euasterids is well-established. Terminal branches longer than 0.8 were again trimmed, resulting in a refined dataset containing 387 orthologs. Final nucleotide alignments were estimated with PRANK v.150803 (Löytynoja, 2014) and cleaned for a minimum of 30% column occupancy using the pxclsq function in phyx. Alignments for the 387 orthologs were concatenated using pxcat in phyx and a maximum likelihood (ML) species tree and rapid bootstrap support (200 replicates) with was inferred using RAxML v8.2.4 with the command raxmlHPC-PTHREADS, the option -f a, and a separate GTRCAT model of evolution estimated for each ortholog; this resulted in a topology we refer to as the maximum likelihood topology (MLT). In order to more fully characterize the likelihood space for these data, 200 regular (i.e. non-rapid) bootstraps with and without the MLT as a starting tree were conducted in RAxML using the -b option in separate runs. Trees for each individual ortholog were also estimated individually using RAxML, with the GTRCAT model of evolution and 200 rapid bootstraps using the option -f a. A coalescent-based maximum quartet support species tree (MQSST) was estimated using ASTRAL v5.6.2 (Zhang et al., 2018) with the resulting 387 ortholog trees. In order to investigate the possible effect of ML tree search algorithm on phylogenetic inference for the 387 ortholog supermatrix, an additional ML tree was estimated using RAxML v8.2.11 and the command raxmlHPC-PTHREADS-AVX and with IQ-TREE v1.6.1 with -m GTR+Γ -n 0 and model partitions for each ortholog specified with the -q option (Nguyen et al., 2015). Likelihood scores of the best scoring tree for each of these tree search algorithms were compared to determine whether the MLT was the topology with the best likelihood score.

### Gene-wise log-likelihood comparisons

A comparison of gene-wise likelihood support for the MLT against a conflicting backbone topology recovered in all 200 rapid bootstrap replicates (i.e. the rapid bootstrap topology, RBT) was conducted using a two-topology analysis (Shen et al., 2017; Walker et al., 2018a). We chose to investigate the RBT topology because even though the MLT received the best likelihood score recovered by any of the tree search algorithms, the MLT was never recovered by RAxML rapid bootstrap replicates, suggesting that the MLT was not broadly supported in likelihood space. A three-topology comparison was also conducted by calculating the gene-wise log-likelihoods of a third topology where the MLT was modified such that the interfamilial backbone relationships were those recovered in the ASTRAL topology (AT); this constructed tree was used in lieu of the actual ASTRAL topology to minimize the effect of intrafamilial topological differences on likelihood calculations. In both tests, the ML score for each gene is calculated while constraining the topology under multiple alternatives. Branch lengths were optimized for each input topology and GTR+Γ was optimized for each supermatrix partition (i.e. each ortholog) for each topology (Shen et al., 2017; Walker et al., 2018a). Results from both comparisons were visualized using a custom R script (R Core Team, 2019). An edge-based analysis was performed to compare the likelihoods of competing topologies while allowing gene tree conflict to exist outside the relationship of interest (Walker et al., 2018a). Our protocol was similar to that of Walker et al. (2018a), except that instead of a defined “TREE SET”, we used constraint trees with clades defining key bipartitions corresponding to each of three competing topologies using the program EdgeTest.py (Walker et al., 2019). The likelihood for each gene was calculated using RAxML-ng with the GTR+Γ model of evolution, using the brlopt nr_safe option and an epsilon value of 1×10^-6^ while a given relationship was constrained (Kozlov et al., 2019). The log-likelihood of each gene was then summed to give a likelihood score for that relationship. An additional edge-based analysis was conducted to investigate the placement of Ebenaceae using the program Phyckle (Smith et al., 2018), which uses a supermatrix and a set of constraint trees specifying conflicting relationships and reports, for each gene, the ML as well as the difference between the best and second-best topology. This allows quantification of gene-wise support at a single edge as well as how strongly the relationship is supported in terms of likelihood.

### Gene duplication comparative analysis

Homolog clusters were filtered such that sequences shorter than half the median length of their cluster and clusters with more than 4000 sequences were removed in order to minimize artifacts due to uncertainty in homolog tree estimation. Homolog trees were then re-estimated with IQ-TREE with the GTR+Γ model and SH-aLRT support followed by cutting of internal branches longer than 1.0 and terminal branches longer than 0.8 inferred substitutions per base pair. Rooted ingroup clades were extracted with the procedure from Yang et al. (2015) with all non-ericalean taxa as outgroups. Gene duplications were inferred with the program phyparts, requiring at least 50% SH-aLRT support in order to avoid the inclusion of very poorly supported would-be duplications (Smith et al., 2015). By mapping gene duplications in this way to competing topological hypotheses (i.e. the MLT, RBT, and AT), as well as several hypothetical topologies employed in order to reveal the number of gene duplications uniquely shared between clades that were never recovered as sister in the species trees, we determined the number of duplications uniquely shared among several ericalean clades.

When using tree-based methods to infer the placement of gene duplications, the location to which duplications are mapped depends on the topology of the phylogeny. Therefore, a gene duplication can appear along a branch in the species tree other than that for which the homolog tree actually contains evidence because duplications are mapped to the bipartition in the species tree that contains *all* the taxa that share a given duplication in the homolog tree. For example, if in a homolog tree, taxon A and taxon B share a gene duplication, but in the species tree taxon C is sister to taxon A, and taxon B sister to those, then the duplication will be mapped to the branch that includes all three taxa, because the duplication is mapped to the smallest clade that includes both taxon A and taxon B. However, if the duplication is instead mapped to a species tree where A and B are sister, the relevant duplication would instead be mapped to the correct, more exclusive branch that specifies the clade for which there is evidence of that duplication in the homolog tree (i.e. only taxa A and B). Once it has been determined how many duplications are actually supported by the homolog trees, comparisons between competing species relationships can be made. It is important to note that incomplete taxon sampling and other biases should be considered when applying such a comparative test of gene duplication number. Assuming there are duplicated genes present in the taxa of interest, clades with a greater number of sampled taxa or more complete transcriptomes will likely share more duplications simply due to the fact that more genes will have been sampled in the dataset. Furthermore, the location to which a duplication is mapped will depend on which assemblies contain transcripts from duplicated genes, whether or not those duplicated genes are present in the genome of any particular organism. Therefore, we investigate the use of gene duplications shared amongst clades as an additional, relative metric of topological support capable of corroborating other results (i.e. that if clades share many gene duplications unique to them, they are more likely to be closely related), while recognizing that the absolute number of duplications shared by various clades are impacted by imperfect sampling.

### Synthesizing support for competing topologies

We reviewed support for each of the three main topological hypotheses (i.e. MLT, RBT and AT) and determined the most commonly supported interfamilial backbone. Because the comparative gene duplication analysis and constraint tree analyses both supported the RBT over other candidates, and the MLT was found to occupy a narrow peak in likelihood space based on bootstrapping and a two-topology test, the RBT was the most commonly supported backbone and was further explored with additional measures of support. Quartet support was assessed on the RBT using the program Quartet Sampling with 1000 replicates (Pease et al., 2018). We used this procedure to measure quartet concordance (QC), quartet differential (QD), quartet informativeness (QI), and quartet fidelity (QF). Briefly, QC measures how often the concordant relationship is recovered with respect to other possible relationships, QD helps identify if a relationship has a dominant alternative, and QI correspond to the ability of the data to resolve a relationship of interest, where a quartet that is at least 2.0 log-likelihood (LL) better than the alternatives is considered informative. Finally, QF aids in identifying rogue taxa (Pease et al., 2018). Gene conflict with the corroborated topology was assessed using the bipartition method as implemented in phyparts using gene trees for the 387 orthologs, which were first rooted on the outgroups *Helianthus annuus* and *Solanum lycopersicum* using a ranked approach with the phyx command pxrr. Gene conflict was assessed both requiring node support (BS≥70) and without any support requirement; the support requirement should help reduce noise in the analysis, but we also ran the analysis without a support requirement to ensure that potentially credible (but only partially-supported) bipartitions were not overlooked. Results for both were visualized using the program phypartspiecharts (https://github.com/mossmatters/phyloscripts/).

### Expanded ortholog datasets

In order to explore the impact of dataset construction, orthologs were inferred from the homolog trees as for the 387 ortholog set but with modifications to taxon requirements and refinement procedures described below. For each dataset, sequences were aligned separately with both MAFFT and PRANK and cleaned for 30% occupancy. A supermatrix was constructed with each and an ML tree was estimated with IQ-TREE with and without a separate GTR+Γ model partition for each ortholog in order to test the effect of model on phylogenetic inference. Individual ortholog trees were estimated with RAxML and used to construct an MQSST with ASTRAL.

#### 2045 ortholog set

Orthologs were filtered such that there was no minimum number of taxa and at least two tips from each of the following five groups: 1) Primulaceae; 2) Polemoniaceae and Fouquieriaceae; 3) Lecythidaceae; 4) outgroups including *Solanum* and *Helianthus* as well as Marcgraviaceae and Balsaminaceae (i.e. the earliest diverging ericalean clade in all previous analyses); and 5) all other taxa. This filtering resulted in a dataset with 2045 orthologs. Ortholog tree support for conflicting placements of Ebenaceae was assessed for the PRANK-aligned orthologs using Phyckle.

#### 4682 ortholog set

Orthologs were not filtered for any taxon requirements. Sequences were aligned with MAFFT, ortholog trees were estimated with RAxML and terminal branches longer than 0.8 were trimmed. Sequences were then realigned separately with MAFFT and PRANK and cleaned as before.

#### 1899, 661, and 449 ortholog sets

To assess the effect of requiring *Helianthus* and *Solanum* outgroups in the 387 ortholog set and to further explore the effect of taxon requirements on the inferred topology, each homolog tree that produced an ortholog in that dataset was re-estimated in IQ-TREE with SH-aLRT support. Calculating this support allowed visual assessment of whether uncertain homolog tree construction was affecting the ortholog identification process. This did not appear to be a major issue as orthologs (i.e. clades of ingroup tips subtended by outgroup tips) within homolog trees typically received strong support as monophyletic.

Following homolog tree re-estimation, orthologs were identified as described above with no minimum taxa requirement. Sequences were aligned with MAFFT and any taxon with >75% missing data for a given ortholog was removed. Filtered alignments were then realigned separately with both MAFFT and PRANK, and cleaned for 30% column occupancy. This resulted in a dataset with 1899 orthologs. To investigate the influence of taxon requirements, two subsets of this 1899 ortholog set were generated by requiring a minimum of 30 and 50 taxa, resulting in 661 and 449 orthologs respectively. Gene tree conflict was assessed for the 449 MAFFT-aligned dataset by rooting all ortholog trees on all taxa in Balsaminaceae and Marcgraviaceae (i.e. the balsaminoid Ericales) since all previous analyses showed this clade to be sister to the rest of Ericales; phyparts was then used to map ortholog tree bipartitions to the ML tree for this dataset and the results were visualized with phypartspiecharts.

### Estimating substitutions supporting contentious clades

In order to gain additional insight into the magnitude of signal present in ortholog alignments informing various relationships, we developed a procedure that identifies clades of interest within an ortholog tree and uses the estimated branch length leading to that clade, multiplied by the length of the corresponding sequence alignment, to estimate the number of substitutions implied by that branch. Applying this approach to the trees from the 449 ortholog MAFFT-aligned dataset with appropriate taxon sampling to allow rooting on a member of the balsaminoid Ericales, we calculated the approximate number of substitutions that implied by the branch leading to the most recent common ancestor (MRCA) of two or more of the following clades: Primulaceae, Polemoniaceae+Fouquieriaceae, Lecythidaceae, Ebenaceae, and Sapotaceae. In addition, we assessed substitution support for several non-controversial relationships, namely that each of the following was monophyletic: Polemoniaceae+Fouquieriaceae, Lecythidaceae, and the non-balsaminoid Ericales. Mean and median values for substitution support were calculated and a distribution of these values was plotted using custom R scripts.

### Synthesis of uncertainty, consensus topology, and genome duplication inference

We took into consideration all previous results, including those of the expanded datasets, gene tree conflict, and substitution support, to determine which relationships were generally well-supported by the results of this study, and which were not. In cases where an interfamilial or intrafamilial relationship remained irresolvable when considering the preponderance of the evidence (i.e. was not supported by plurality of methods employed after accounting for nested, conflicting relationships), that relationship was not included in the consensus topology. Gene duplications were mapped to the consensus topology using the methods of the comparative duplication analysis described above. Internodes on inferred species trees with notably high numbers of gene duplications were used as one line of evidence for assessing putative WGDs. Further investigation into possible genome duplications was conducted by plotting the number of synonymous substitutions (K_s_) between paralogs according to the methods of Yang et al. (2015) after removing sequences shorter than half the median length of their cluster. Multispecies K_s_ values (i.e. ortholog divergence K_s_ values) for selected combinations of taxa were generated according to the methods of Wang et al. (2018). The effect of evolutionary rate-heterogeneity among ericalean species was investigated by conducting a multispecies K_s_ analysis of each non-balsaminoid ericalean taxon against *Impatiens balsamifera*, since all evidence suggests each of these species pairs have the same MCRA (i.e. the deepest node in the Ericales phylogeny).

Because each pair has the same MRCA, the resulting ortholog peak in each case, represents the same speciation event and differences in the location of this peak among K_s_ plots are the result of differences in evolutionary rate among species. In rare cases were the location of the ortholog peak was ambiguous (i.e. these were two or more local maxima near the global maximum) the plot was not considered in the rate-heterogeneity analysis. All single and multispecies K_s_ plots were generated using custom R scripts.

## RESULTS

### Initial 387 ortholog dataset

The concatenated supermatrix for the 387 ortholog dataset contained 441,819 aligned sites with 76.1% ortholog occupancy and 57.8% matrix occupancy. The MLT recovered by RAxML v8.2.4 is shown in Figure 1. The 200 rapid bootstrap trees all contained the same backbone topology (i.e. the RBT) that differed from that of the MLT by one relationship. However, regardless of which tree search algorithm was used, the likelihood score for the MLT was better than for any other topology recovered for this dataset. This indicates that while the MLT is only supported by a narrow peak in likelihood space, it was indeed the topology with the best likelihood based on the methods employed for the 387 ortholog dataset. The MLT contained the clade Polemoniaceae+Fouquieriaceae as sister to a clade consisting of Ebenaceae, Sapotaceae, and what is referred to here as the “Core” Ericales: the clade that includes Actinidiaceae, Diapensiaceae, Ericaceae, Pentaphylacaceae, Roridulaceae, Sarraceniaceae, Styracaceae, Symplocaceae, and Theaceae. The RBT instead placed Lecythidaceae sister to Ebenaceae, Sapotaceae, and the Core (a hypothetical clade herein referred to as ESC), which was recovered by 100% of rapid bootstrap replicates (Fig. 1). A gene-wise log-likelihood analysis comparing these two topologies is shown in Appendix S3. The cumulative log-likelihood difference between the MLT and RBT was approximately 3.57 in favor of the MLT; there were 27 orthologs that supported the MLT over the RBT by a score larger than this and the exclusion of any of these from the supermatrix could cause the RBT to become the topology with the best likelihood. Both of these topologies were considered as candidates in a search for a corroborated interfamilial backbone. Regular (i.e. non-rapid) bootstrapping in RAxML resulted in 6.5% support for Polemoniaceae+Fouquieriaceae sister to ESC (i.e. the MLT) unless the MLT was given as a starting tree, under which conditions the MLT topology received 100% bootstrap support. Unguided, regular bootstrapping instead suggested strong support (90%) for a clade consisting of Lecythidaceae and ESC (Appendix S4). Interfamilial relationships recovered in the ASTRAL topology (AT) were congruent with those of the MLT except that Primulaceae was recovered as sister to Lecythidaceae+ESC, but with only 54% local posterior probability (Appendix S5). A total of seven intrafamilial relationships differed between the AT and MLT. The AT was considered as a third candidate for an interfamilial backbone.

**Figure 1.**
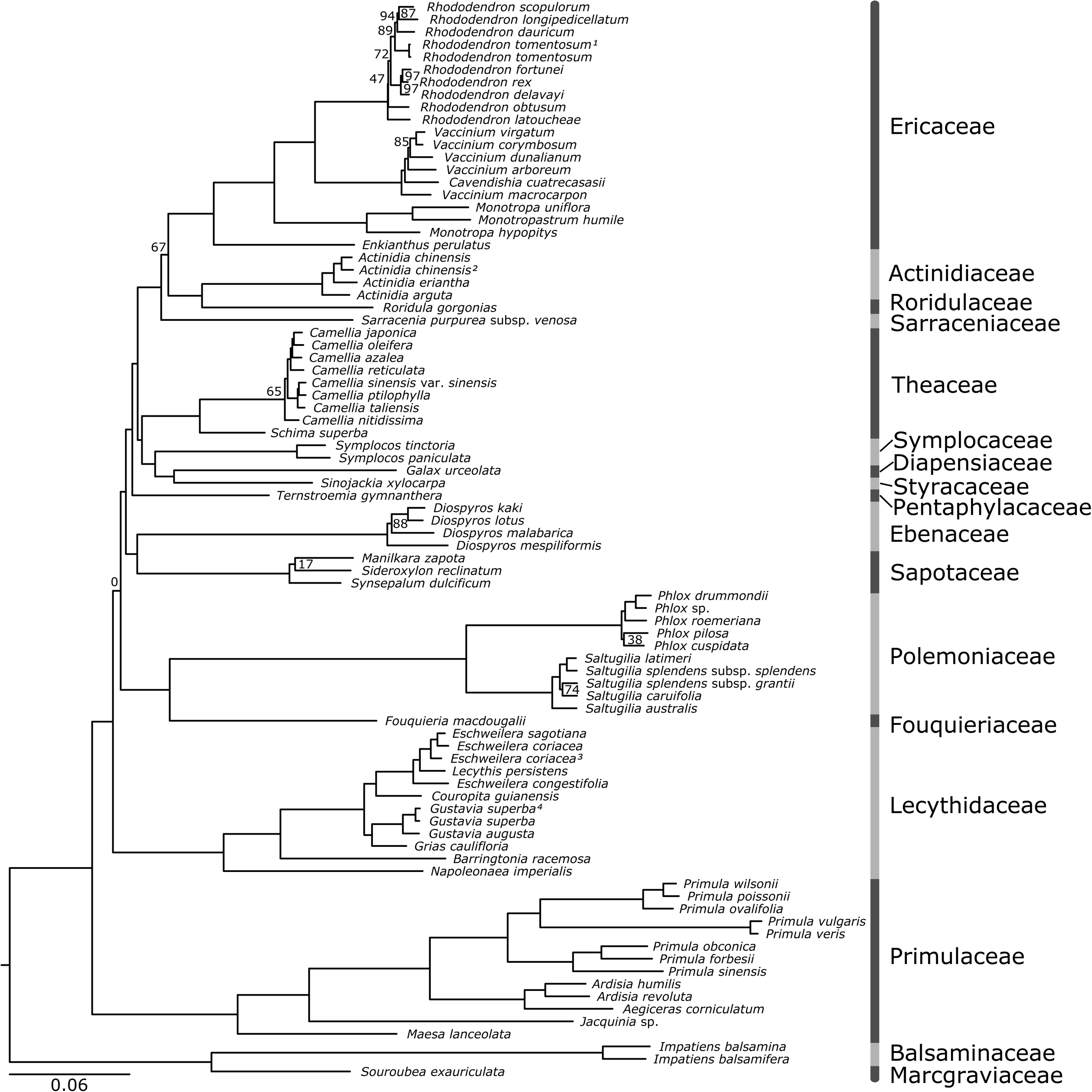
Maximum likelihood topology recovered for a 387 ortholog dataset using RAxML. Nodes receiving less than 100% rapid BS support are labeled. Branch lengths are substitutions per base pair. The node that determined the placement of Lecythidaceae received zero support (i.e. the ML topology was never recovered by a rapid bootstrap replicate).

There was an effect of the ML tree search algorithm that may be important to note for future phylogenomic studies (Zhou et al., 2017). Re-conducting an ML tree search for the 387 ortholog supermatrix with RAxML v8.2.11 using PTHREADS-AVX architecture and -m GTRCAT or with IQ-TREE and GTR+Γ resulted in a species tree with a different topology than under any other conditions for the 387 ortholog dataset. The final log-likelihood of the original ML tree was −5948732.774. Using the PTHREADS-AVX architecture returned a final likelihood of −5948752.263, while the best log-likelihood reported by IQ-TREE was −5949077.382 (19.489 and 344.607 log-likelihood points worse than the RAxML v8.2.4 results, respectively).

### Gene-wise log-likelihood comparative analysis across three candidate topologies

The results from the gene-wise comparisons of likelihood contributions showed that there were 155 orthologs in the dataset that most strongly supported the MLT, 90 supported the RBT, and 142 supported the AT (Appendix S6). The AT had a cumulative log-likelihood score that was >300 points worse, even though more individual genes support the AT over the RBT (Appendix S6).

### Edge-based comparative analyses across three candidate topologies

Of the three candidate topologies investigated by constraining key edges, Lecythidaceae sister to ESC received the best score, while Primulaceae sister to ESC received the worst (Appendix S7). Similarly, the clade Lecythidaceae+Polemoniaceae+Fouquieriaceae+ESC received a better likelihood score than the clade Lecythidaceae+Primulaceae+ESC. Regarding the placement of Ebenaceae for this dataset, Ebenaceae+Sapotaceae was supported by the highest number of orthologs whether or not two log-likelihood support difference was required (Appendix S8). However, a number of orthologs support each of the investigated placements for Ebenaceae and less than half of genes supported any placement over the next best alternative by at least two log-likelihood points.

### Gene duplication comparative analysis

Gene duplications mapped to each candidate backbone topology and the five additional hypothetical topologies revealed differing numbers of shared duplications that can be used as a metric of support among candidate topologies (Fig. 2). Regarding which clade is better supported as sister to ESC, Lecythidaceae uniquely shareed 433 duplications with ESC, more than twice as many as either alternative. Polemoniaceae+Fouquieriaceae shared 1300 unique duplications with Lecythidaceae+ESC, more than Primulaceae which shared 626 unique duplications (Fig. 2).

**Figure 2.**
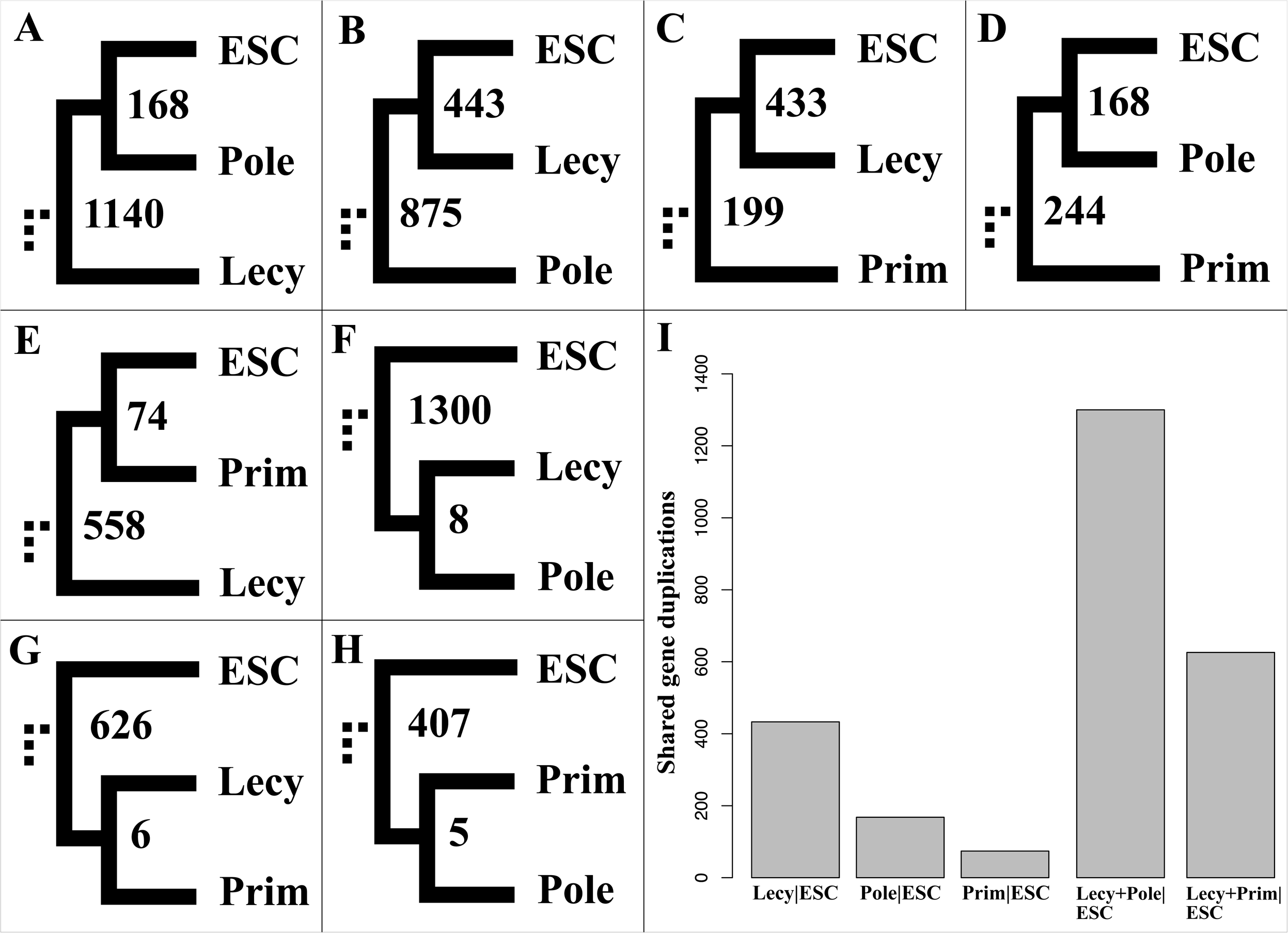
(A-H) Gene duplications with at least 50% SH-aRLT support in homolog trees mapped to several topologies recovered from the 387 ortholog dataset including the maximum likelihood topology (A) the rapid bootstrap topology (B) the ASTRAL topology (C) and several hypothetical topologies that demonstrate evidence for shared duplications in clades not recovered with species tree methods (D-H). Names of clades are abbreviated to four letters, ESC represents the clade Ebenaceae+Sapotaceae+Core Ericales, and “Pole” represents the clade Polemoniaceae+Fouquieriaceae in all cases. I) Bar chart showing the number of uniquely shared gene duplications between clades that can be considered a metric of support for distinguishing among conflicting topological relationships.

### A corroborated topology for the 387 ortholog set

The above comparative analyses supported one topology among the three candidates identified for the 387 ortholog dataset, namely the RBT (Fig. 2; Appendix S7-9). In this tree, all taxonomic families are recovered as monophyletic. Marcgraviaceae and Balsaminaceae (i.e. the balsaminoid Ericales) are sister, and form a clade that is sister to the rest of Ericales (i.e. the non-balsaminoid Ericales). Ebenaceae and Sapotaceae form a clade that is sister to the Core. The monogeneric Fouquieriaceae is sister to Polemoniaceae. A clade containing Symplocaceae, Diapensiaceae, and Styracaceae is sister to Theaceae. Roridulaceae is sister to Actinidiaceae and these together form a clade that is sister to Ericaceae. A grade containing Primulaceae, Polemoniaceae and Fouquieriaceae, and Lecythidaceae leading to ESC is supported by the rapid bootstrapping, comparative duplication, and constraint-tree analyses.

### Quartet sampling

In our results and discussion we consider a QC score of (≥0.5) to be strong support as this signifies strong concordance among quartets (Pease et al., 2018). We found varying levels of support for several key relationships in the RBT (Appendix S9). The monophyly for all families received strong support (QC ≥ 0.90). There was strong support (QC=0.54) for the node placing Lecythidaceae sister to ESC, while equivocal support (QC=0.035) for Polemoniaceae+Fouquieriaceae sister to Lecythidaceae and ESC. Within ESC, there was moderate support (QC=0.28) for Ebenaceae sister to Sapotaceae, but poor (QC=-0.13) support for this clade as sister to the Core. There was strong support (QC=0.85) for the clade including Symplocaceae, Diapensiaceae, and Styracaceae and moderate support (QC=0.26) for this clade as sister to Theaceae. Roridulaceae was very strongly supported (QC=0.99) as sister to Actinidiaceae but there was no support for this clade as sister to Ericaceae (QC=-0.23). However, the monophyly of the clade that includes Roridulaceae, Actinidiaceae, Ericaceae, and Sarraceniaceae received very strong support (QC=0.90).

The QD scores for several contentious relationships indicate that discordant quartets tended to be highly skewed towards one conflicting topology as indicated by scores below 0.3 (Pease et al., 2018). However, the QD score for the relationship placing Lecythidaceae sister to ESC in the RBT was 0.53, indicating relative equality in occurrence frequency of discordant topologies. Similarly, the relationship placing Polemoniaceae+Fouquieriaceae sister to Lecythidaceae+ESC received a QD score of 0.83, indicating that among alternative topologies (e.g. Primulaceae sister to the clade Lecythidaceae+ESC), there was no clear alternative to the RBT recovered through quartet sampling for this dataset. The QI scores for all nodes defining interfamilial relationships were above 0.9, indicating that in the vast majority of sampled quartets there was a tree that was at least two log-likelihoods better than the alternatives. The QF scores for all but one taxon were above 0.70 and the majority were above 0.85, which shows that rogue individual taxa were not a major issue (Pease et al., 2018).

### Conflict analyses

Assessing ortholog tree concordance and conflict for the 387 ortholog set mapped to the RBT showed that backbone nodes that differed between candidate topologies were poorly supported with the majority of orthologs failing to achieve ≥70% bootstrap support (Appendix S10). Among informative orthologs, the majority of trees conflict with any candidate topology at these nodes and there is no dominant alternative to the RBT. There were 77 ortholog trees with appropriate taxon sampling that placed Primulaceae sister to the rest of the non-balsaminoid Ericales and 37 did so with at least 70% bootstrap support. The clade Lecythidaceae+Primulaceae+ESC was recovered in 47 ortholog trees, and in 17 with at least 70% bootstrap support. Of the 33 ortholog trees that contained the clade Primulaceae+Ebenaceae, nine did so with at least 70% bootstrap support.

### Expanded ortholog sets

Seven combinations of relationships along the backbone were recovered in analyses of the expanded ortholog sets which we term E-I though E-VII for reference (Fig. 3). Among these, the ML tree estimated from the 2045-ortholog PRANK-aligned, partitioned supermatrix placed Lecythidaceae and Ebenaceae in a clade sister to Sapotaceae and the Core (E-II), with the other interfamilial relationships recapitulating those of the RBT; the topology recovered by ASTRAL for these orthologs (E-VII) placed Ebenaceae, Lecythidaceae, Polemoniaceae+Fouquieriaceae, and Primulaceae as successively sister to Sapotaceae and the Core. In regard to the placement of Ebenaceae for the 2045 PRANK-aligned set, the edge-based Phyckle analysis showed that 432 of orthologs in this dataset with appropriate taxon sampling supported Ebenaceae sister to Primulaceae, while 202 did so by at least two log-likelihood over any alternative (Appendix S11). However, 416 orthologs supported Ebenaceae sister to Sapotaceae (158 with ≥2LL), and 316 supported Ebenaceae sister to Lecythidaceae (140 with ≥ 2LL). When the 2045 ortholog set was aligned with MAFFT to produce a supermatrix, the ≥ resulting ML topology (E-I) was such that the clade Primulaceae+Ebenaceae were sister to Sapotaceae, with that clade sister to the Core and Polemoniaceae+Fouquieriaceae sister to all of those. When ASTRAL was run with the 2045 MAFFT-aligned orthologs, the topology recovered (E-III) placed Polemoniaceae+Fouqieriaceae sister to the Core, with the clade Ebenaceae+Primulaceae and Lecythidaceae sucessively sister to those.

**Figure 3.**
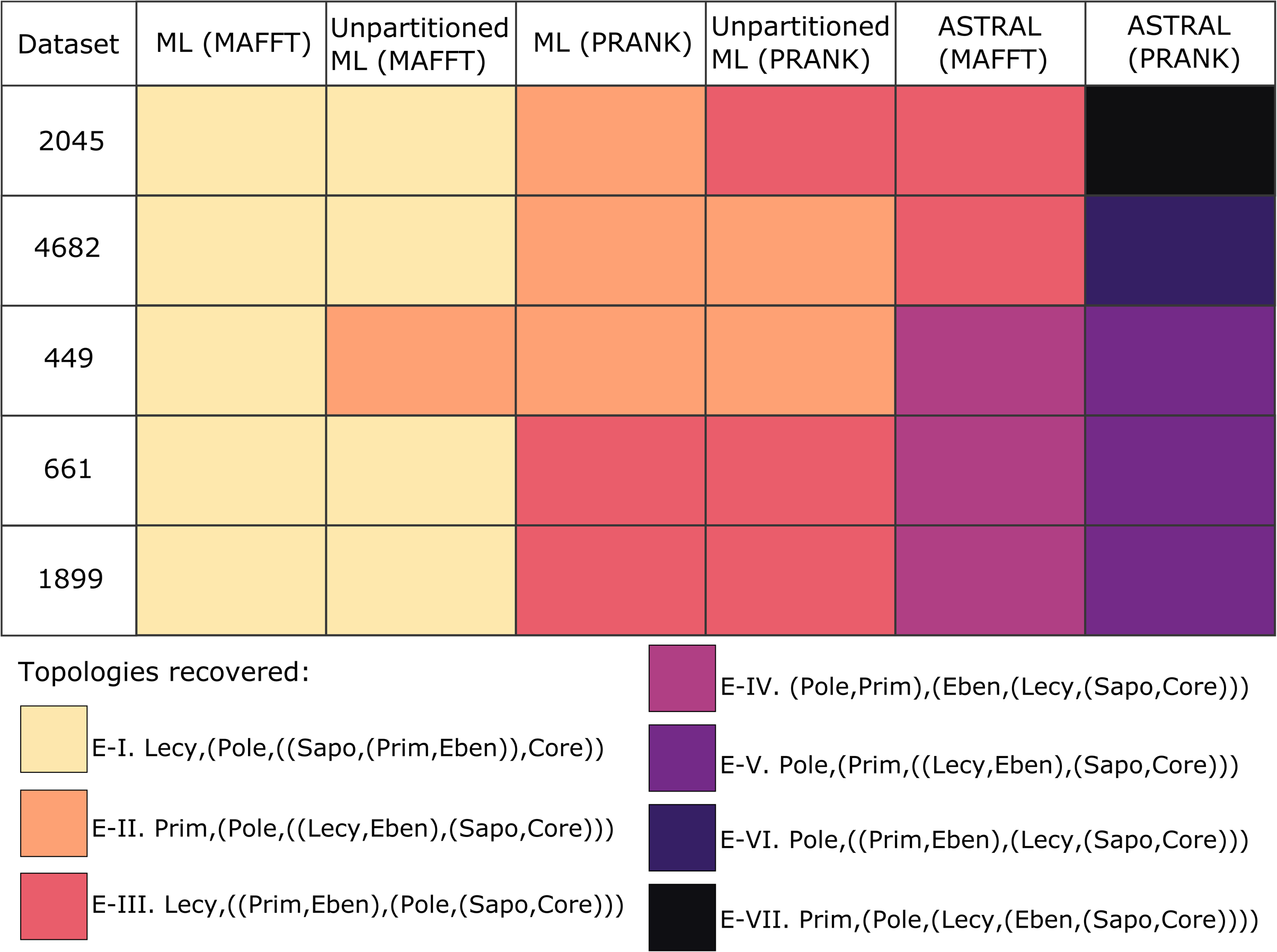
Topologies recovered from several combinations of ortholog datasets and species tree methods explored as alternatives to the focal 387 ortholog dataset. Each color corresponds to a unique backbone topology recovered in these analyses. Names of clades are abbreviated to four letters and “Pole” represents the clade Polemoniaceae+Fouquieriaceae in all cases.

The backbone topology resulting from the 4682 ortholog PRANK-aligned, partitioned supermatix was the same as that recovered with the 2045 ortholog and the same methods (E-II). The ASTRAL topology for the 4682 ortholog PRANK-aligned dataset (E-VI) placed Lecythidaceae sister to Sapotaceae and the Core, with Primulaceae+Ebenaceae and Polemoniaceae+Fouquieriaceae successively sister to those. When the 449, 661, or 1899 ortholog set were aligned with MAFFT to produce a supermatrix, the resulting ML backbone topology was the same as that of the MAFFT-aligned 2045 and 4682 ortholog sets (E-I), except when the 449 ortholog supermatrix was run without partitioning (E-II). When ortholog alignments were produced with PRANK, the ML backbone recovered from the 449 and ortholog set was the same as that of the 2045 PRANK-aligned ML tree (E-II). The backbone of the ML trees produced from the 1899 and 661 PRANK-aligned ortholog sets were the same as that of the 2045 ortholog MAFFT-aligned ASTRAL tree (E-III). The 440, 661, and 1899 PRANK-aligned ortholog sets all produced the same backbone topology in ASTRAL (E-V). The backbone topologies recovered by ASTRAL for the 449, 661, and 1899 MAFFT-aligned ortholog sets also agree with one another (E-VI), but conflict with all other species trees recovered in this study.

### Synthesis of uncertainty and determination of an overall consensus

In the species trees generated with the 387, 449, 661, 1899, 2045, and 4682 ortholog datasets, the relationships among several taxonomic families were in conflict with one another (Fig. 3). Many of these relationships also conflict with the thoroughly investigated RBT, which was shown to be very well-supported by the 387 ortholog set. Nine of ten ML trees generated with MAFFT-aligned supermatrices in the expanded datasets recovered Primulaceae and Ebenaceae sister to one another and forming a clade with Sapotaceae (E-I), however this pattern was not recovered under any other circumstances (Fig. 3). In all, Primulaceae was recovered as sister to Ebenaceae in 17 of the 33 species trees generated in this study (51.5%), but these families were sister in only eight of the 24 trees (33.3%) where they did not form a clade with Sapotaceae. Sapotaceae was recovered as sister to the Core in 21 of the 33 species trees (63.6%). Thus, there is no majority consensus that reconciles these conflicting relationships; Primulaceae is only sister to Ebenaceae in a majority of trees if those that also contain the clade Sapotaceae+Ebenaceae+Primulaceae are considered and that topology is in direct conflict with Sapotaceae+Core, a relationship that is recovered in a majority of trees. In addition, edge-wise support for a sister relationship between Primulaceae and Ebenaceae was dataset dependent and showed equivocal gene-wise support. The positions of Lecythidaceae or the clade containing Polemoniaceae and Fouquieriaceae that is not agreed by a majority of species trees. Given this, the relationships among Primulaceae, Polemoniaceae and Fouqieriaceae, Lecythidaceae, Ebenaceae, and Sapotaceae are not resolved in the consensus topology. All other interfamilial relationships were unanimously supported by all species trees except for the relationships between the clade Symplocaceae, Diapensiaceae, and Styracaceae sister to Theaceae, which was recovered in 29 of the 33 species trees (87.9%).

### Gene and genome duplication inference

Gene duplications mapped to the consensus topology shows that the largest number of duplications occurs on the branch leading the non-Balsaminoid Ericales (Fig. 4). Notable numbers of gene duplications also appear along branches leading to *Actinidia* (Actinidiaceae), *Camellia* (Theaceae), several members of Primulaceae, *Rhododendron* (Ericaceae), *Impatiens* (Balsaminaceae), and Polemoniaceae (i.e. *Phlox* and *Saltugilia* in this study). If gene duplications are mapped to the single most-commonly recovered species tree in this study (E-1), 3485 duplications occur along the branch leading to the non-balsaminoid Ericales (Appendix S12). The K_s_ plots show peaks for several of the recent duplications, evidenced by peaks between 0.1 and 0.5 (Appendix S13). The K_s_ plots for most, but not all, taxa also contain a peak that appears between 0.8 and 1.5. The multispecies K_s_ plots for some ericalean taxa paired with a member of Cornales show an ortholog peak with a higher K_s_ value (i.e. farther to the right) than the paralog peaks near 1.0, suggesting two separate duplication events that each occurred after the divergence of the two orders. Some other combinations of ingroup and outgroup taxa resulted in an ortholog peak to the left of the paralogs peaks (Appendix S14). When comparing the position of unambiguous ortholog peaks of all non-balsaminoid ericalean taxa to *Impatiens balsamifera*, the K_s_ value of the peak varied between values of 1.01 to 1.57 for *Schima superba* (Theaceae) and *Primula poissonii* (Primulaceae) respectively, indicating substantial rate heterogeneity in the accumulation of synonymous substitutions among ericalean taxa and implying that in the most extreme cases, some species have accumulated synonymous substitutions 55% faster than others (Appendix S15).

**Figure 4.**
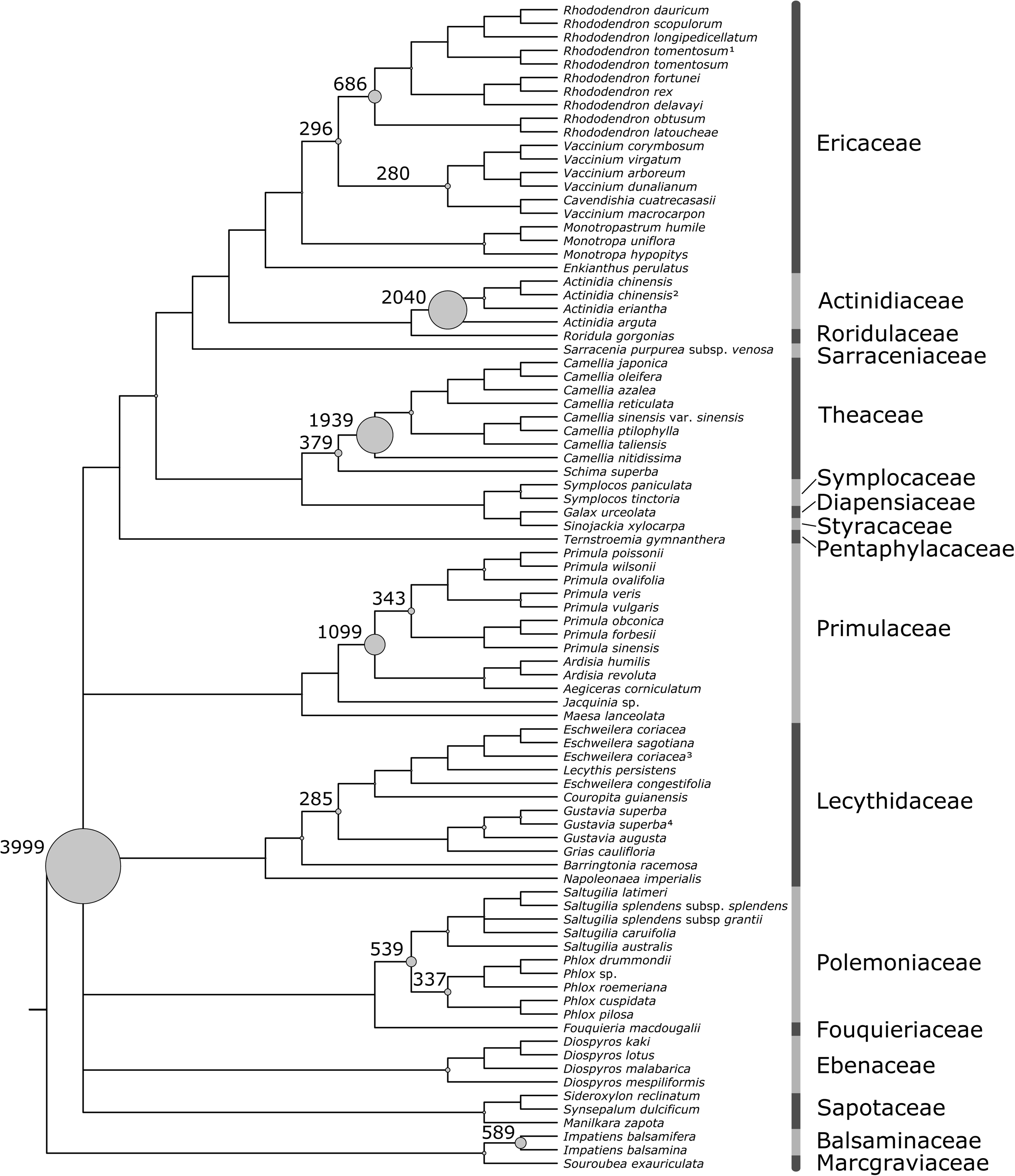
Gene duplications mapped to a cladogram of the consensus topology. Contentious relationships not supported by a plurality of methods were collapsed to a polytomy. The diameter of circles corresponds to the number of inferred duplications, and cases with at least 250 are labelled along branches. The single largest number of duplications occurs on the branch leading the non-Balsaminoid Ericales. Verified genome duplications in the Theaceae and the Actinidiaceae appear as the second and third largest numbers of inferred gene duplications respectively.

### Estimating substitutions supporting contentious clades

The estimated number of substitutions informing clades containing at least two families whose backbone placement is contentious tended to be much less than for relationships that garnered widespread support. These contentious clades had a median value of 8.65 estimated substitutions informing them, compared to 30.08, 75.09, and 144.00 informing the monophyly of Polemoniaceae and Fouqieriaceae, Lecythidaceae, and the non-Balsaminoid Ericales, respectively, in the MAFFT-aligned ortholog trees (Fig. 5). In trees for the same orthologs with alignments generated instead with PRANK, the corresponding median values were 8.96, 30.73, 74.16 and 145.98 respectively.

**Figure 5.**
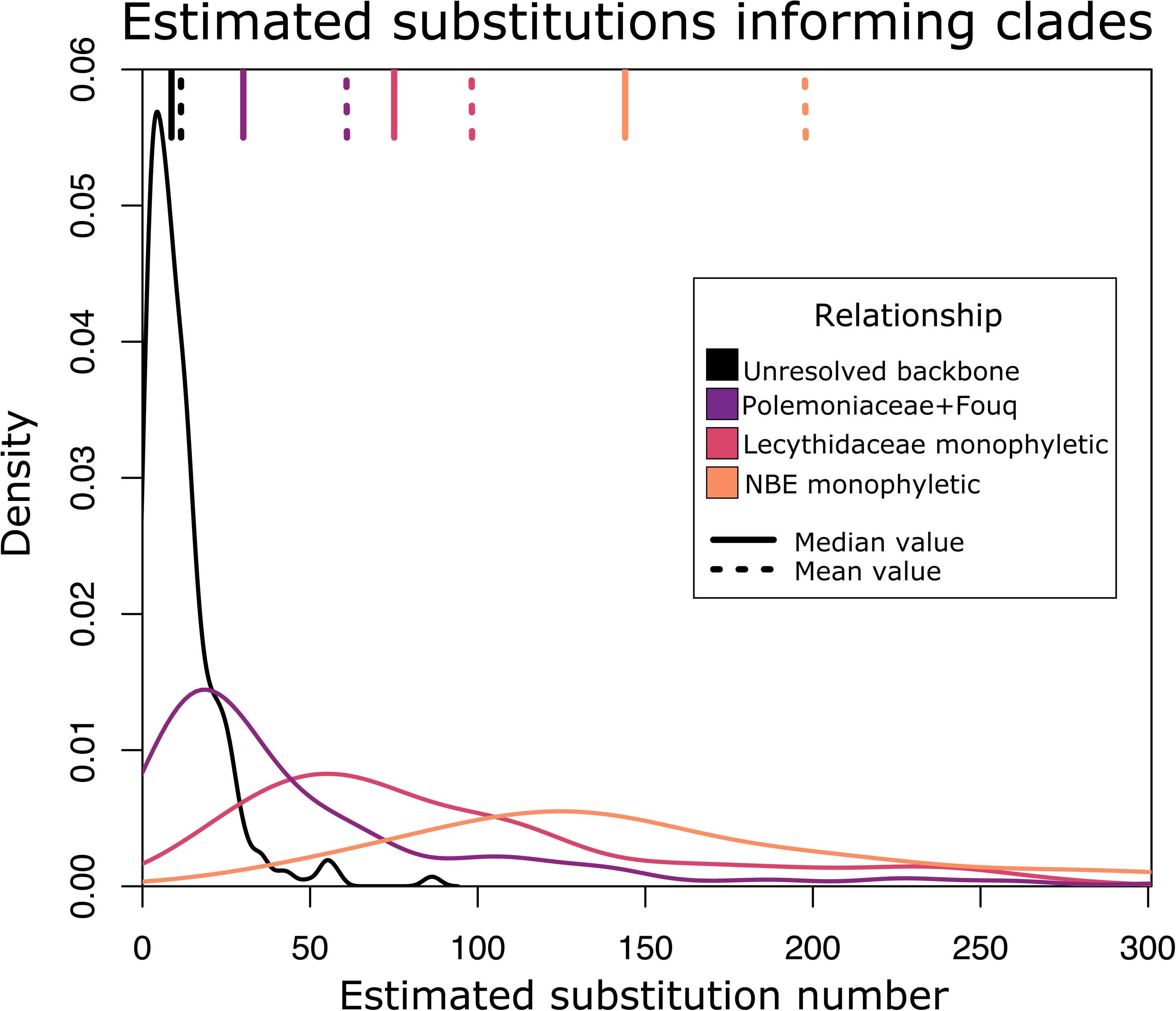
Estimated number of substitutions supporting clades using rooted orthologs from the 449 ortholog MAFFT-aligned dataset. The median number of estimated substitutions was 8.65 for clades that would resolve the polytomy in the consensus topology, compared to 30.08, 75.09, and 144.00 informing the monophyly of Polemoniaceae and Fouqieriaceae, Lecythidaceae, and the non-Balsaminoid Ericales respectively.

## DISCUSSION

Our focal dataset, consisting of 387 orthologs, supports an evolutionary history of Ericales that is largely consistent with previous work on the clade for many, but not all, relationships (Appendix S9). The balsaminoid clade, which includes the families Balsaminaceae, Tetrameristaceae (not sampled in this study), and Marcgraviaceae was confidently recovered as sister to the rest of the order as has been shown previously (e.g. Geuten et al., 2004; Rose et al., 2018; Gitzendanner et al., 2018). Similarly, the monogeneric family Fouquieriaceae is sister to Polemoniaceae, the para-carnivorous Roridulaceae are sister to Actinidiaceae and the circumscription of Primulaceae sensu lato is monophyletic (Rose et al., 2018). The majority of analyses for the 387 ortholog dataset recovered a sister relationship between Sapotaceae and Ebenaceae, with that clade sister to the Core Ericales and Lecythidaceae sister to those. Notably, the topology supported suggests that Primulaceae diverged earlier than has been recovered in most previous phylogenetic studies and does not form a clade with Sapotaceae and Ebenaceae as has been suggested by others, including Rose et al. (2018). The topology recovered by Gitzendanner et al. (2018) using coding sequences from the chloroplast placed Primulaceae sister to Ebenaceae, with those as sister to the rest of the non-basaminoid Ericales, though that study did not include sampling from Lecythidaceae. In addition to the traditional phylogenetic reconstruction methods, by applying the available data on gene duplications as a metric of support and leveraging methods that make use of additional phylogenetic information present in the supermatrix, we were able to more holistically summarize the evidence present in a 387 ortholog dataset in an effort to resolve the Ericales phylogeny (Figs. 1-2; Appendix S3-S10).

It has been shown repeatedly that large phylogenetic datasets have a tendency to resolve relationships with strong support, even if the inferred topology is incorrect (Seo, 2008). However, some of our results suggest extreme sensitivity to tree-building methods. For example, the initial ML analysis resulted in an ML topology (MLT) with zero rapid bootstrap support for the placement of Lecythidaceae (Fig. 1). The rapid bootstrap consensus instead unanimously supported Lecythidaceae sister to a clade including Ebenaceae, Sapotaceae and the Core Ericales (i.e. the RBT). Gene-wise investigation of likelihood contribution confirmed that these two topologies have very similar likelihoods but did not identify outlier genes that seemed to have an outsized effect on ML calculation (Appendix S3). Instead, the cumulative likelihood influence of the 387 genes in the supermatrix provides nearly equal support for the two topologies, while ASTRAL resulted in a third, and regular bootstrapping recovered an even more diverse set of topologies (Appendix S4). These results suggest that there are several topologies with similar likelihood scores for this dataset. Despite the fact that the additional comparative analyses applied to the 387 ortholog dataset supported a single alternative among those investigated (i.e. the RBT), the recovery of multiple topologies by various tree-building and bootstrapping methods suggested that the criteria used to generate and filter orthologs could have marked potential to influence the outcome of our efforts to resolve relationships amongst the families of Ericales.

In addition to sensitivities associated with tree building methods, we investigated additional datasets constructed with a variety of methods and filtering parameters to shed further light on the nature of the problem of resolving the ericalean phylogeny. While it is clear from investigation of gene tree topologies for the 387 ortholog dataset that phylogenetic conflict is the rule rather than the exception, the expanded datasets show that this is true for a variety of approaches to dataset construction and not simply an artifact of one approach. While the monophyly of most major clades and the relationships discussed above were recovered across these datasets, we also demonstrate that many combinations of contentious backbone relationships can be recovered depending on the methods used in dataset construction, alignment, and analysis (Fig 3).

### An unresolved consensus topology

Based on the data available, we suggest that while the relationships recovered in the Core Ericales and within most families are robust across methodological alternatives, there is insufficient evidence to resolve several early-diverging relationships along the ericalean backbone. We, therefore, suggest that the appropriate representation, until further data collection efforts and analyses show otherwise, is as a polytomy (Fig. 4). Whether this is biological or the result of data limitations remains to be determined. A biological polytomy (i.e. hard polytomy) can be the result of three or more populations diverging rapidly without sufficient time for the accumulation of nucleotide substitutions or other genomic events to reconstruct the patterns of lineage divergence. Most of the inferred orthologs contain little information useful for inferring relationships along the backbone of the phylogeny; we investigated this explicitly by estimating the number of nucleotide substitutions that inform these backbone relationships and find that branches that would resolve the polytomy were based on a small fraction of the number of substitutions that informed better-supported clades (Fig. 5). The phylogenetic signal presented in this study results in extensive gene tree conflict, albeit mostly with low support (Appendix S11). The major clades of Ericales may or may not have diverged simultaneously; however, if divergence occurred rapidly enough as to preclude the evolution of genomic synapomorphies, then a polytomy may be a reasonable representation of such historical events rather than signifying a shortcoming in methodology or taxon sampling.

The high levels of gene tree conflict and lack of a clear consensus among datasets for a resolved topology is likely to have multiple causes. Among these is the fact that this series of divergence events seems to have happened relatively rapidly over the course of around 10 million years (Rose et al., 2018). We also find evidence that a WGD is likely to have occurred before or during this ancient radiation (Fig. 4; Appendix S12). If this is the case, differential gene loss and retention during the process of diploidization is likely to complicate our ability to resolve the order of lineage divergences. In addition, we cannot exclude the possibility of hybridization and introgression between these ancient lineages at some point in the past, as hybridization has been documented between plants that have been diverged for tens of millions of years (e.g. Arias et al., 2014; Rothfels et al., 2015). It is even possible that some of the lineages involved in such introgression have gone extinct in the intervening 100 million years and that introgression from such now-extinct “ghost lineages” could represent a potentially insurmountable obstacle to fully understanding the events that lead to the diversity of forms we now find in Ericales. We chose not to test explicitly for evidence of hybridization here, due to the seeming equivocal phylogenetic signal present in most gene trees for these contentious relationships. We suggest that interpreting the generally weak signal present in most conflicting gene trees as anything other than a lack of reliable information, runs a high risk of overinterpreting these data since network analyses and tests for introgression generally treat gene tree topologies as fixed states known without error. However, future studies could potentially find such an approach to be appropriate for explaining the high levels of conflict among orthologs, but should carefully consider alternative explanations for gene tree discordance.

### Gene and genome duplications in Ericales

Results from gene duplication analyses showed evidence for several whole genome duplications in Ericales, including at least two, in *Camellia* and *Actinidia*, that have been verified using sequenced genomes (Fig. 6; Huang et al., 2013; Xia et al., 2017; Shi et al., 2010). Our results strongly support the conclusion drawn by Wei et al., (2018) that the most recent WGD in *Camellia* is distinct from the *Ad*-α WGD that occurred in Actinidiaceae after that clades diverged from other Ericales. We propose the name *Cm*-α for this WGD, which is shared by all *Camellia* in our study and may or may not also be shared by *Schimia*, the only other genus in Theaceae that we sampled. Future studies with broader taxon sampling should be able determine whether the *Cm*-α WGD is shared by other genera in Theaceae or if it is exclusive to *Camellia*.

**Figure 6.**
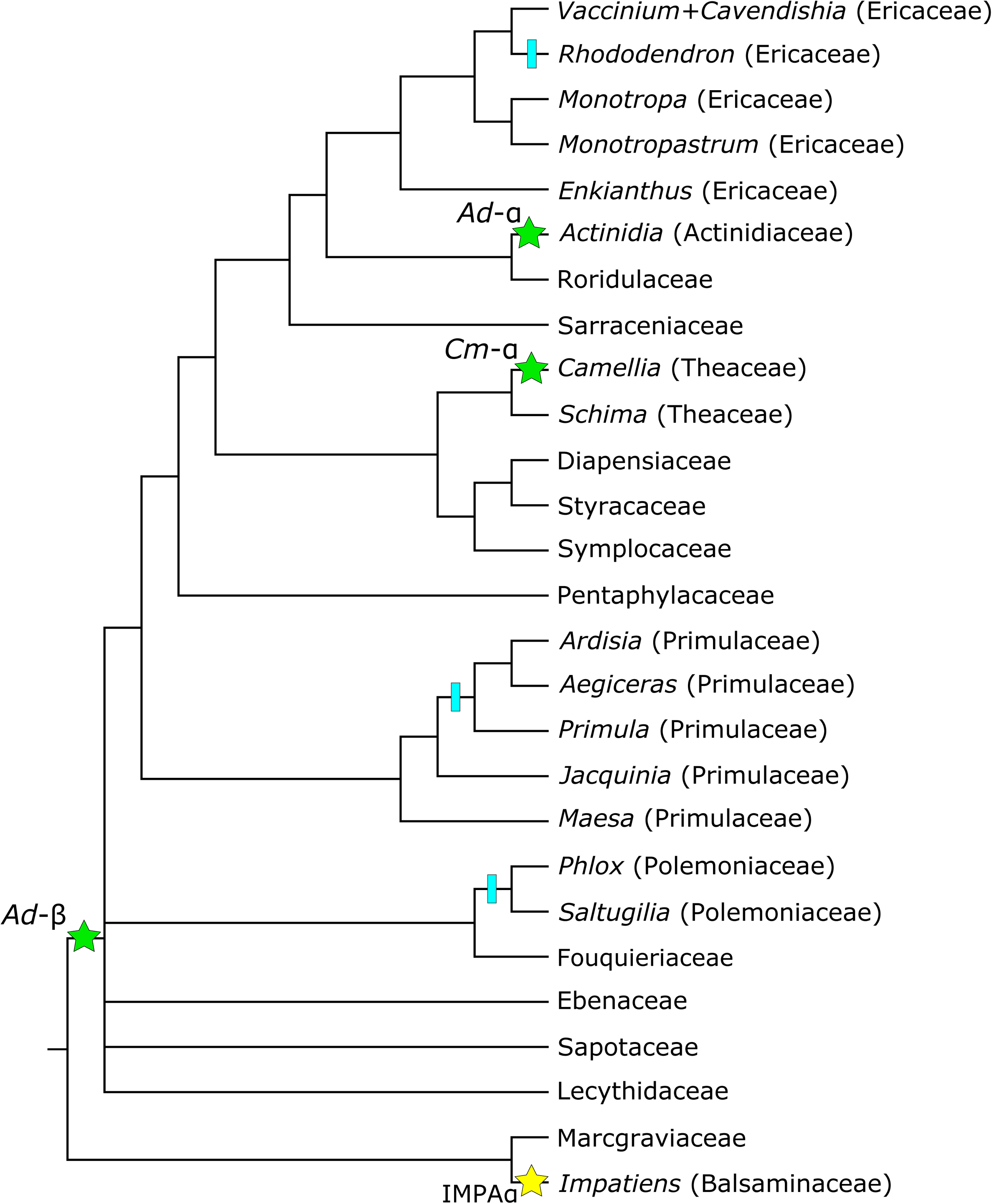
Placements of putative whole genome duplications (WGDs) on a consensus phylogeny of Ericales. Green stars represent WDGs that have been corroborated by sequenced genomes (Shi. et al., 2010; Xia et al., 2017; Soza et al., 2019). Yellow stars represent WGDs that have been proposed previously based on transcriptomes and are corroborated in this study (Leebens-Mack et al., 2019). Blue tick marks identify branches with evidence of previously uncharacterized WGDs that should be investigated in future studies. Ad- and Ad-are named β after the WGD first detected in *Actinidia* (Shi et al. 2010). Cm-α is a name proposed in this study α for a WGD unique to Theaceae that has been characterized previously and corroborated here (e.g. Wei et al., 2018). In cases were a possible or confirmed WGD was inferred along a branch leading to or within a botanical family, tips represent the genera sampled in this study. If no lineage-specific WGD was inferred for a family, the tip represents all taxa sampled for that family.

Our inferred genome duplications are concordant with several, but not all, of the conclusions drawn by Leebens-Mack et al. (2019), whose transcriptome assemblies comprised 24 of the 86 ingroup samples for this study. We find evidence for their ACCHα (i.e. *Ad*-α; Shi et al., 2010) and IMPA α WGDs in Actinidiaceae and Balsaminaceae respectively, though our broader taxon sampling additionally reveals that both of these WGDs are shared by multiple species of their respective genera (Fig. 4; Appendix S12). Our results do not support their placement of ACCHβ, which would appear as a WGD shared exclusively by the Core Ericales in this study (Fig. 4; Appendix S12), nor do our results support the existence of DIOSα Leebens-Mack et al. (2019) infer to have occurred along a branch leading to a clade consisting of Polemoniaceae, Fouquieriaceae, Primulaceae, Sapotaceae, and Ebenaceae: a clade never recovered in this study, and recovered in only one of the three species tree methods employed by Leebens-Mack et al. (2019). We do not find evidence for MOUNα in Monotropoidiae and the in-paralog trimming procedure we employed precludes us from addressing the putative SOURα WGD because our sampling includes only one taxon from Marcgraviaceae (Leebens-Mack et al., 2019).

Our results suggest that a WGD occurred along the backbone of Ericales, either before or after the divergence of the balsaminoid clade, but after Ericales diverged with Cornales (Fig. 4; Appendix S12). Given the extent of the topological uncertainty recovered along the backbone of Ericales in this and other studies (e.g. Fig. 3; Gitzendanner et al., 2018; Rose et al., 2018; Leebens-Mack et al., 2019;) and the fact that our single most-commonly recovered backbone (i.e. Topology E-I, Fig. 3; Appendix S12) would imply that most gene duplications occurred along the branch leading to the non-balsaminoid Ericales, we suggest that a single, shared WGD is the most justifiable explanation for the observed data, rather than a more complex series of nested WGD or near-simultaneous WGDs in sister lineages. We also infer notably high numbers of gene duplications along the branches within Ericaceae, Primulaceae, and Polemoniaceae, which suggests that these clades should be further investigated for evidence of novel, lineage-specific whole genome or other major chromosomal duplications (Fig. 4; Fig. 6).

The K_s_ plots for our most ingroup species appear to share two peaks, in addition to peaks corresponding to several lineage specific WGDs (Appendix S13-15). One shared peak occurs between 2.0-2.5 in most taxa, which is often interpreted as to corresponding to the genome duplication shared by all angiosperms (Jiao et al., 2012, Leebens-Mack et al., 2019). Our results show that this peak between 2.0 and 2.5 also includes to the “γ” palaeopolyploidization shared by the core Eudicots (Appendix S13; Jiao et al., 2011). We are able to infer this by evaluating K_s_ plots for *Helianthus*, *Solanum*, and *Beta* (Appendix S13). Because in-paralogs were trimmed in our homolog trees for all taxa, any WGDs in *Helianthus*, *Solanum*, or *Beta* not shared by another taxon in our study (i.e. Ericales and Cornales) will not appear in the K_s_ plot for that species. Therefore, the K_s_ plots for *Helianthus*, *Solanum*, and *Beta* will exclusively display evidence of polyploidization events that occurred before the MRCA of Asterales and Solanales in the cases of *Helianthus* and *Solanum*, or the MRCA of Asterales+Solanales and Caryophyllales in the case of *Beta* (Appendix S13). Leebens-Mack et al. (2019) show that only polyploidizations at least as old as γ should be shared by these taxa and since none have a K_s_ peak with a value less than 2.0, that peak must include the γ event. Our characterization of the γ event is compatible with the conclusions of Qiao et al. (2019), who analyzed 141 sequenced genomes and found that the γ palaeopolyploidization corresponded to a K_s_ peak that ranged between 1.91 to 3.64 for 16 species that have not experienced a WGD since γ. Qiao et al. (2019) also fitted Ks distributions for their taxa with Gaussian mixture models; for *Actinidia chinensis*, fitted Ks peaks occurred at 0.317, 1.016, and 2.415, which correspond respectively to the first (*Ad*-α), second (*Ad*-β), and third (γ) most recent WGDs in *Actinidia*.

Many of our ingroup species share a K_s_ peak occurring between 0.8 and 1.5 (Appendix S13-S14) that seems to correspond to a WGD shared by all non-balsaminoid Ericales (Fig. 4; Appendix S12). We suggest this is the *Ad*-β WGD characterized by Shi et al. (2010) and corroborated by Soza et al. (2019) and Qiao et al. (2019), the ACCHβ WGD recovered by Leebens-Mack et al. (2019), and the genome duplication Wei et al. (2018) concluded was shared between *Camellia* and *Actinidia*. Our study is the first with the necessary taxon sampling to show that the *Ad*-β WGD likely occurred in the ancestor of all or nearly all ericalean taxa and is likely shared by a clade that minimally includes Lecythidaceae, Polemoniaceae, Fouquieriaceae, Primulaceae, Ebenaceae, Sapotaceae, and the Core Ericales. The tree-based methods employed here precluded us from explicitly inferring the number of gene duplications that occurred directly before the divergence of the balsaminoid Ericales, because we employed a procedure that treated all non-ericalean taxa as outgroups for identifying duplicated ingroup clades in the homolog trees. Studies with broader taxonomic foci should investigate whether the balsaminoid clade share the *Ad*-β WGD with the rest of Ericales. Future sampling of transcriptomes and genomes will likely lead to the discovery of additional, lineage-specific WGDs in Ericales and refine our understanding of which taxa share these and other duplication events (Yang et al., 2018).

Our results strongly suggest that uncertainty should be considered when inferring duplications with tree-based methods since the species tree topology can determine where gene duplications appear to have occurred (Fig. 2; Appendix S12; Zwaenepoel and Van de Peer, 2019). The use of K_s_ plots as a second source of information may not completely ameliorate issues caused by topological uncertainty, since K_s_ plots are generally interpreted in the context of an accepted phylogeny (i.e. an error-free phylogeny where well-supported and contentious nodes are treated the same). Furthermore, K_s_ plots are affected by heterogeneity in evolutionary rate, with faster-evolving taxa accumulating synonymous substitutions at faster rates than more slowly evolving lineages, adding an additional complicating factor when comparing K_s_ peaks and synonymous ortholog divergence values across species (Appendix S15; Smith and Donoghue, 2008, Qiao et al., 2019). Technical challenges such as missing data resulting from incomplete transcriptome sequencing, failure to assemble all paralogs in all gene families, biases in taxon sampling, as well as phylogenetic uncertainty in homolog trees, influences where many individual gene duplications appear in this and other studies of non-model organisms and caution should be taken to avoid overinterpreting noisy signal as biological information.

## CONCLUSIONS

The first transcriptomic dataset broadly spanning Ericales and constructed from publicly available data resolves many of the relationships within the clade and supports several relationships that have been proposed previously. Our results confirm genome duplications in Actinidiaceae and Theaceae and provide a more precise placement of a whole genome duplication in an early ancestor of Ericales. We find evidence to suggest additional WGDs in Balsaminaceae, Ericaceae, Polemoniaceae, and Primulaceae. While our results were largely concordant within taxonomic families, the topological resolution of the deep divergences in Ericales are less decisive. We demonstrate that, with the available data, there is not enough information to strongly support any resolution, despite previous studies having considered these relationships resolved. Additional data will be needed to investigate the early divergences of the Ericales. Our analyses demonstrate that uncertainty needs to be thoroughly investigated in phylotranscriptomic datasets, as strong support can be given by different methods for conflicting topologies that can in turn impact the placement of WGDs on phylogenies. Even in a dataset containing hundreds of genes and hundreds of thousands of characters, the criteria used in dataset construction altered the inferred topology. Our results suggest that phylogenomic studies should employ a range of methodologies and support metrics so that topological uncertainty can be more fully explored and reported. The high prevalence of conflict among datasets and the lack of clear consensus in regards to the relationships among several major ericalean clades led us to conclude that a single, fully resolved tree is not supported by these transcriptome data, though we acknowledge that future improvements in sampling might justify their resolution.

## Data Accessibility Statement

Data, full output for analyses, and custom software used in this study will be available on Dryad: DOI Forthcoming.

## Acknowledgements

The authors thank Christopher Dick, Greg Stull, Hannah Marx, Ning Wang, Caroline Edwards, and members of the Smith lab for their comments on the manuscript. Support came from the National Science Foundation; D.A.L and J.F.W. were supported by NSF DEB 1207915, O.M.V. was supported by NSF FESD 1338694, and S.A.S. was supported by NSF DBI 1458466.

## Author Contributions

The study was conceived by D.A.L. and S.A.S. Dataset construction was led by D.A.L. with contributions by O.M.V. and input from J.F.W. Analyses were led by D.A.L. and J.F.W with contributions and input by S.A.S. D.A.L. wrote the manuscript and produced the figures with contributions and input from O.M.V, J.F.W., and S.A.S. All authors read and approved the final draft of the manuscript.

## SUPPLEMENTAL MATERIALS

**Supplemental Appendix 1.** The origins of transcriptome assemblies, reference genomes, and raw reads used in homolog clustering. Citations associated with raw reads available on the National Center for Biotechnology Information Sequence Read Archive are included where such information was provided in the accession record or could be confidently identified through a Google Scholar query of the relevant accession information.

**Supplemental Appendix 2.**
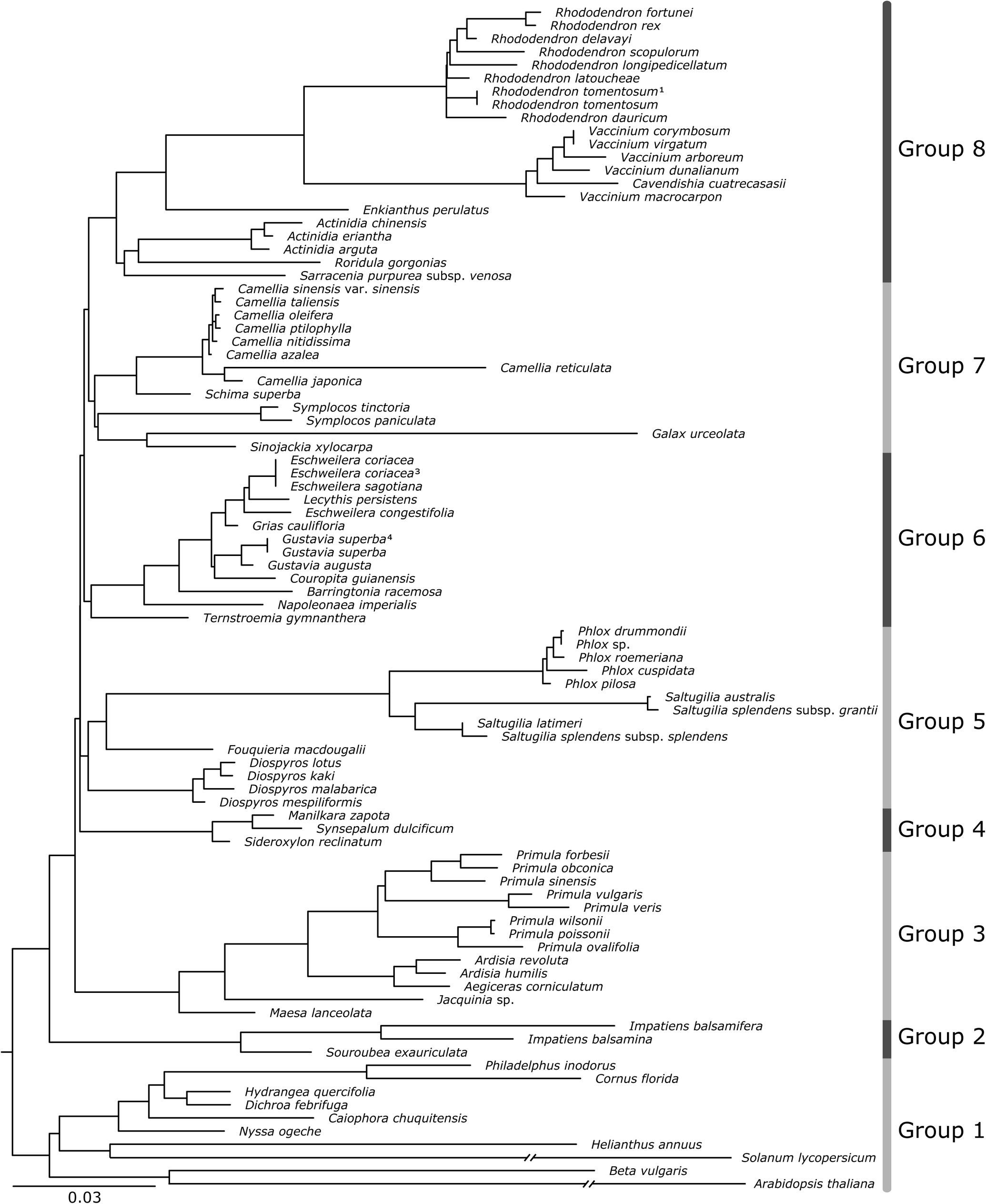
Phylogram inferred using maximum likelihood on a supermatrix consisting of the genes *rpoC2*, *rbcL*, *nhdF*, and *matK*. The topology of the tree was used to assign taxa into eight groups for hierarchical homolog clustering, such that each clustering group was monophyletic and not prohibitively large.

**Supplemental Appendix 3.**
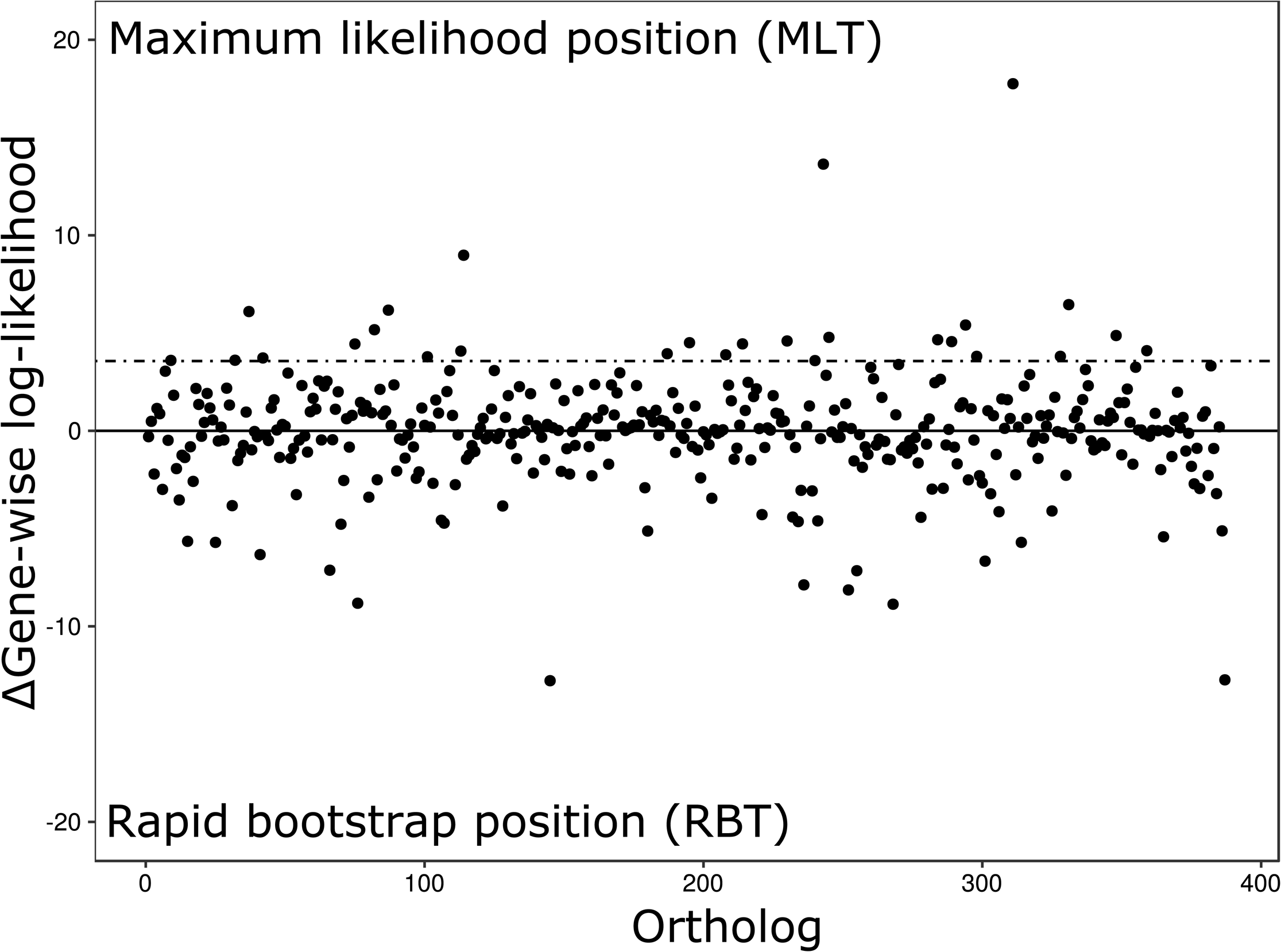
Results of a two-topology test comparing gene-wise support in the 387 ortholog dataset for two alternative placements of Lecythidaceae recovered by the maximum likelihood (ML) search. Delta gene-wise log-likelihood represents the extent to which an ortholog supports the ML topology over that recovered unanimously in rapid bootstraps. The dashed line represents the cumulative difference in log-likelihood between the two competing topologies, such that removing any of the 27 orthologs with a score more positive than that would cause the ML topology to change, even in the absence of obvious outlier genes.

**Supplemental Appendix 4.**
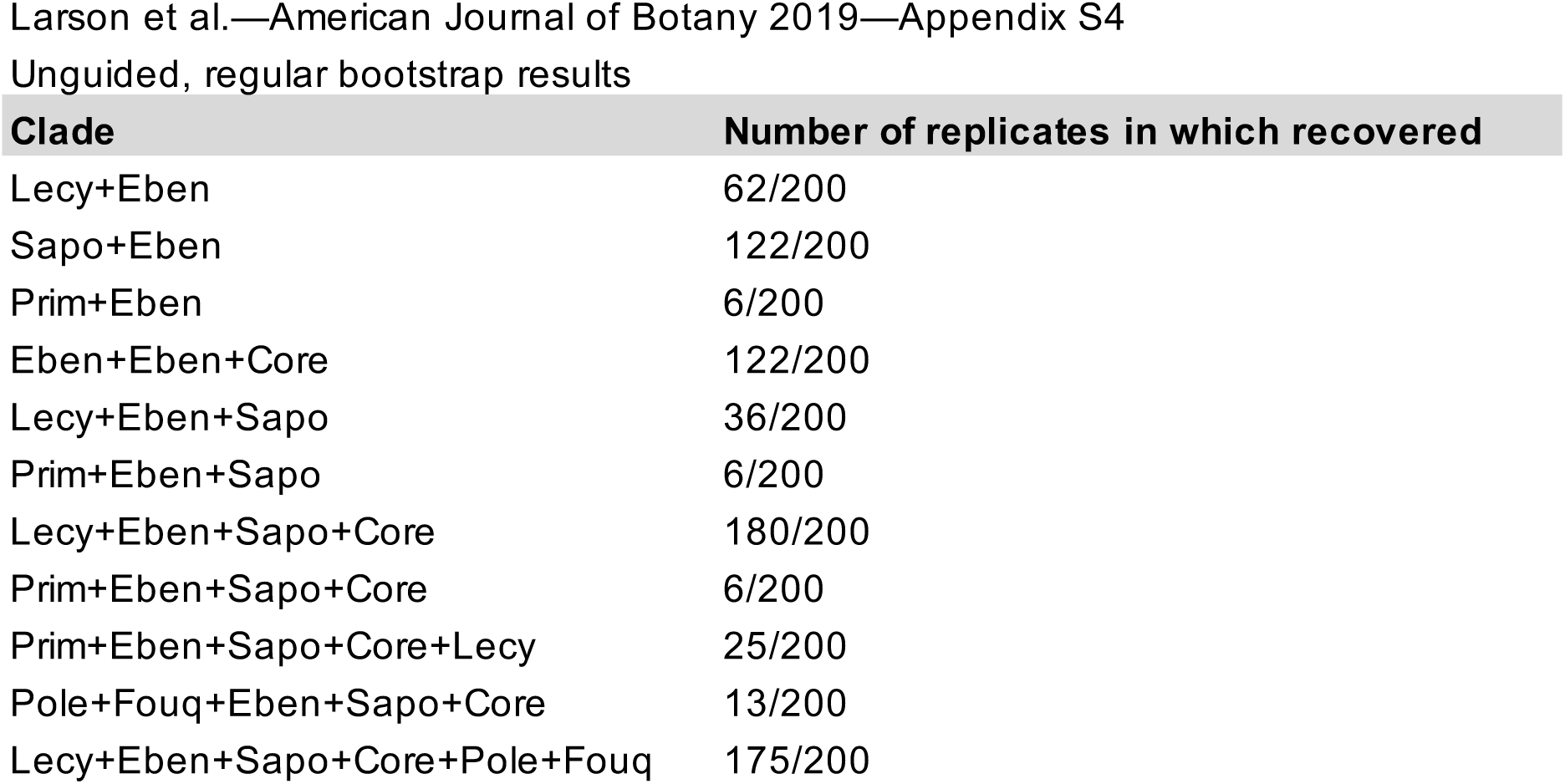
Results of unguided, full bootstrapping with the 387 ortholog supermatrix in RAxML. The number of bootstrap replicates in which various contentious backbone relationships were recovered are reported.

**Supplemental Appendix 5.**
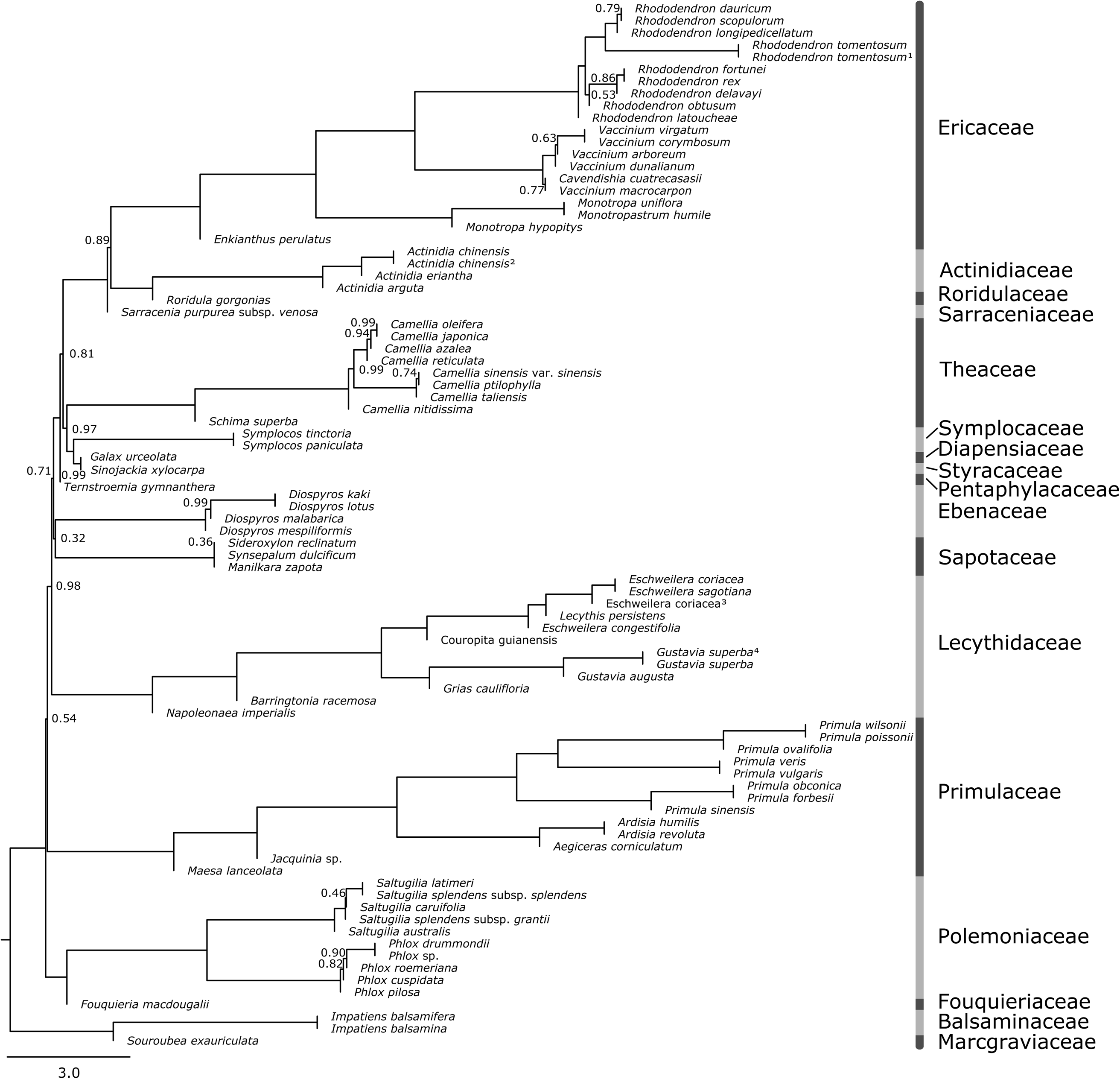
The maximum quartet support species tree topology for the 387 ortholog set generated with ASTRAL. Node support values are ASTRAL local posterior probabilities and nodes receiving support less than 1.0 are labeled. Branch lengths are in coalescent units.

**Supplemental Appendix 6.**
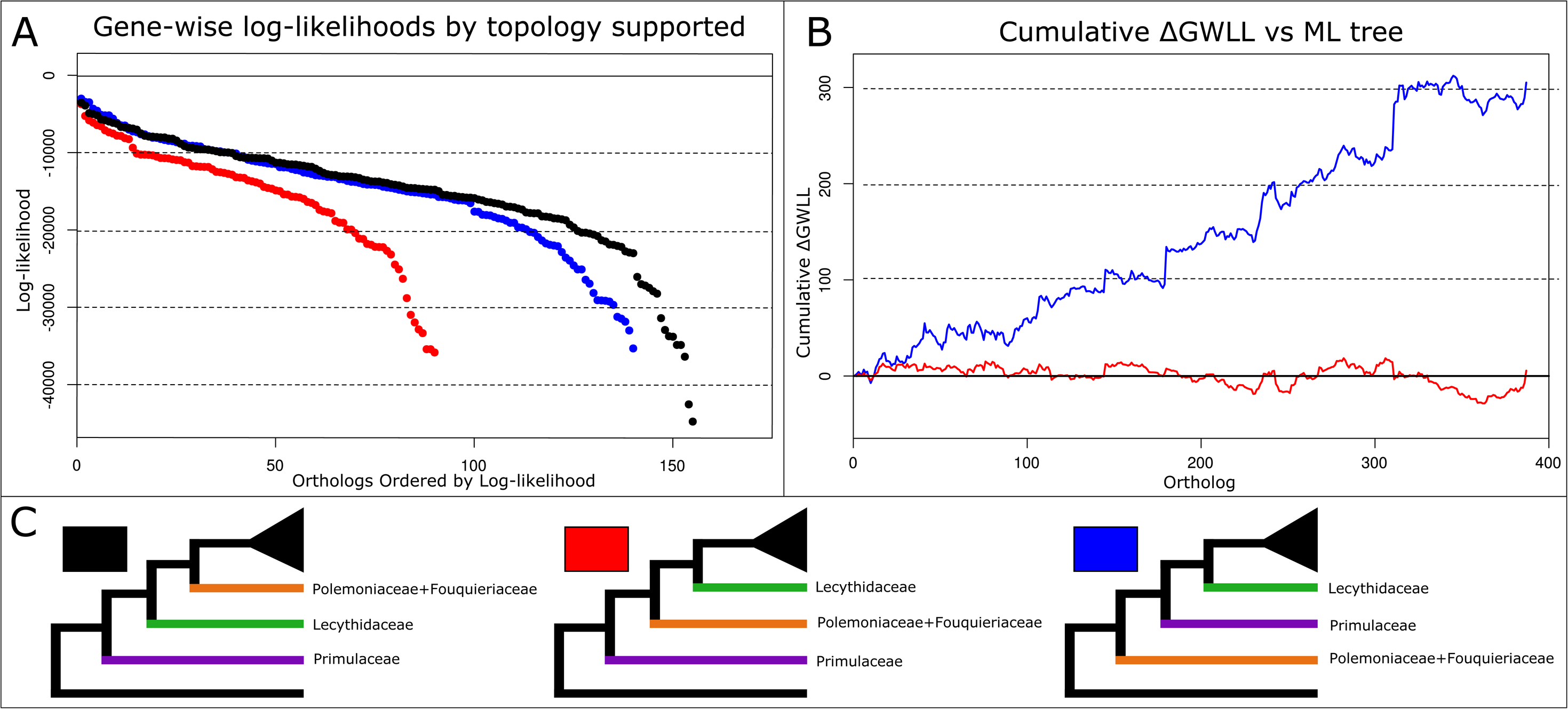
Results from the gene-wise comparisons of likelihood contributions for the three candidate topologies for the 387 ortholog dataset. A) The absolute log-likelihood for each ortholog, sorted by which topology they best support and organized in descending order of likelihood. B) The cumulative difference in log-likelihood relative to the ML topology as orthologs are added across the supermatrix. Delta gene-wise log-likelihood (ΔGWLL) represents the extent to which the topology has a worse score than the ML topology; values below zero indicate that a topology has a better likelihood than the ML topology. C) Schematic of the three candidate topologies in question.

**Supplemental Appendix 7.**
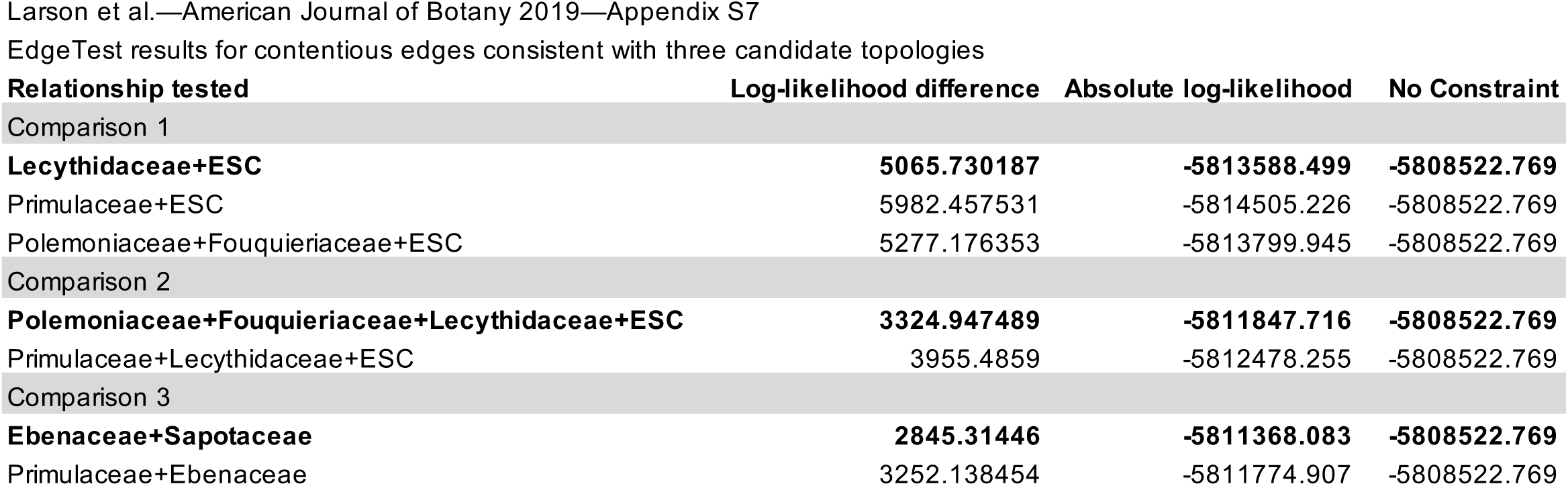
Log-likelihood penalties incurred by constraining contentious edges consistent with each of the three candidate topologies for a 387 ortholog dataset using EdgeTest. Direct comparisons of scores between conflicting relationships can be used as a metric of support with lower scores suggesting stronger support.

**Supplemental Appendix 8.**
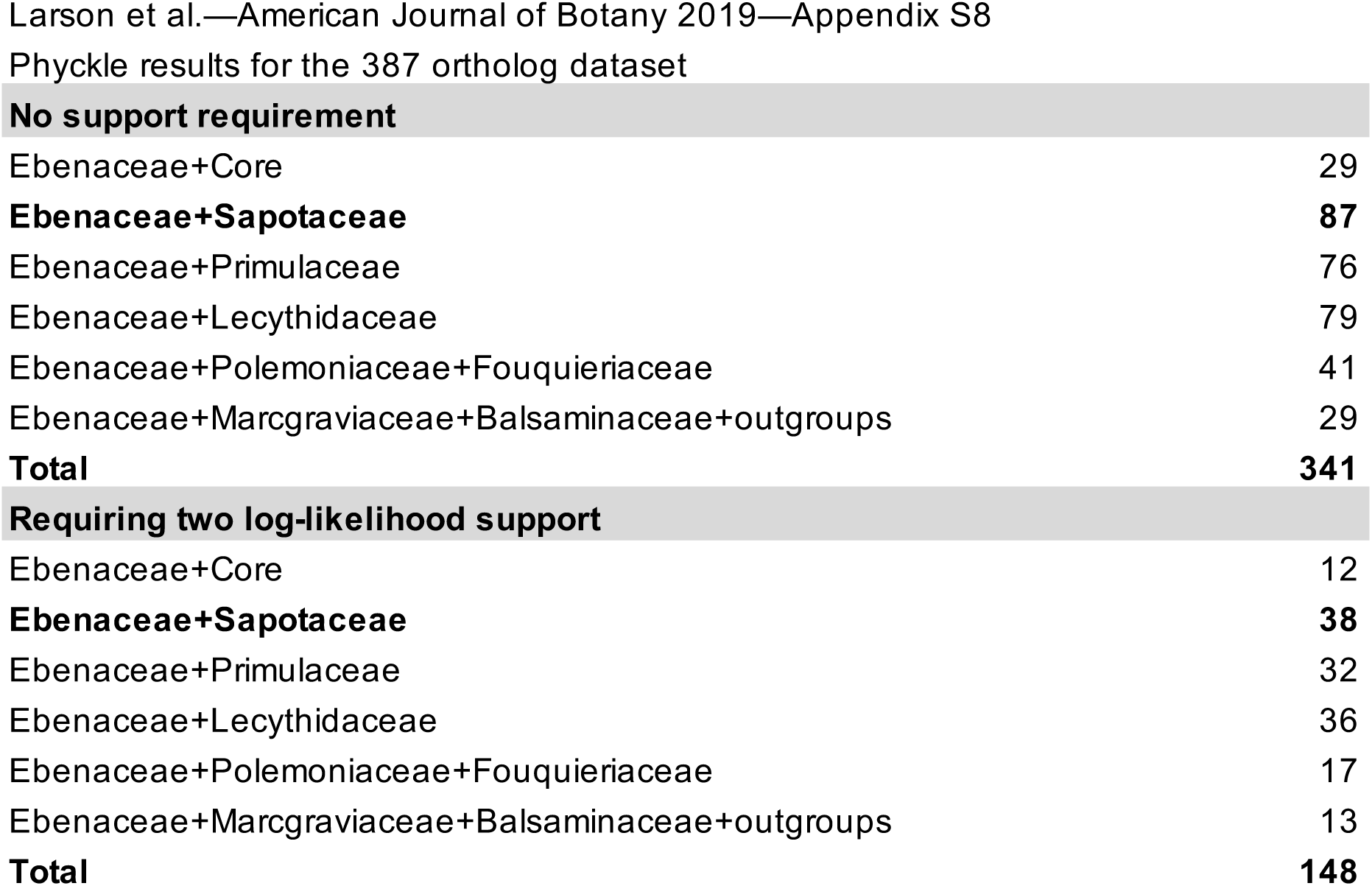
Phyckle results regarding the placement of Ebenaceae for the 387 ortholog dataset. Of the relationships examined, Ebenaceae+Sapotaceae was supported by the highest number of orthologs whether or not two log-likelihood support difference was required.

**Supplemental Appendix 9.**
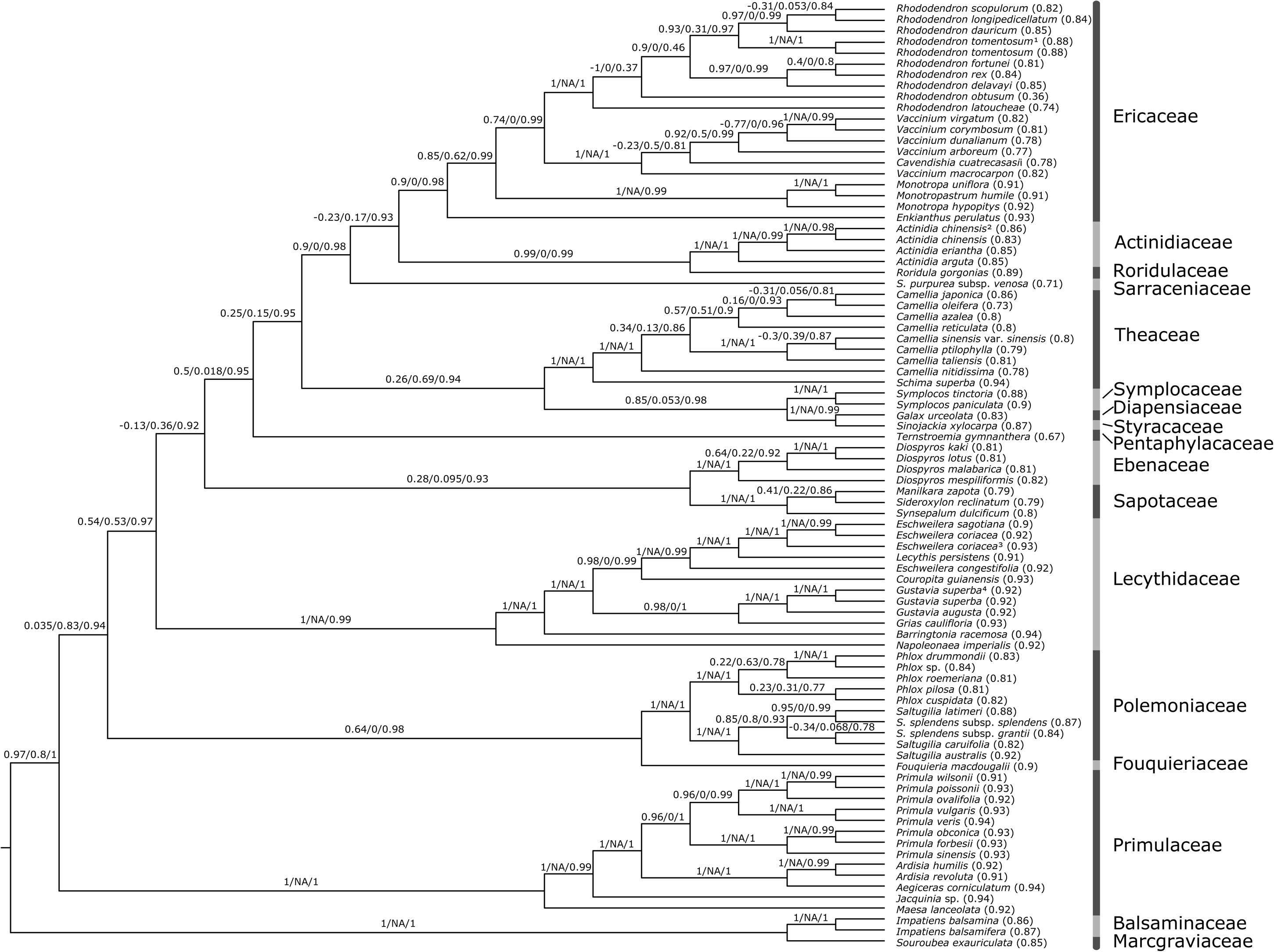
Cladogram showing results from quartet sampling on the rapid bootstrap topology (RBT) from a 387 ortholog dataset. Branch labels show quartet concordance (QC), quartet differential (QD), and quartet informativeness (QI) respectively for each relationship. Quartet fidelity (QF) for each taxon is shown in parentheses after the relevant taxon label.

**Supplemental Appendix 10.**
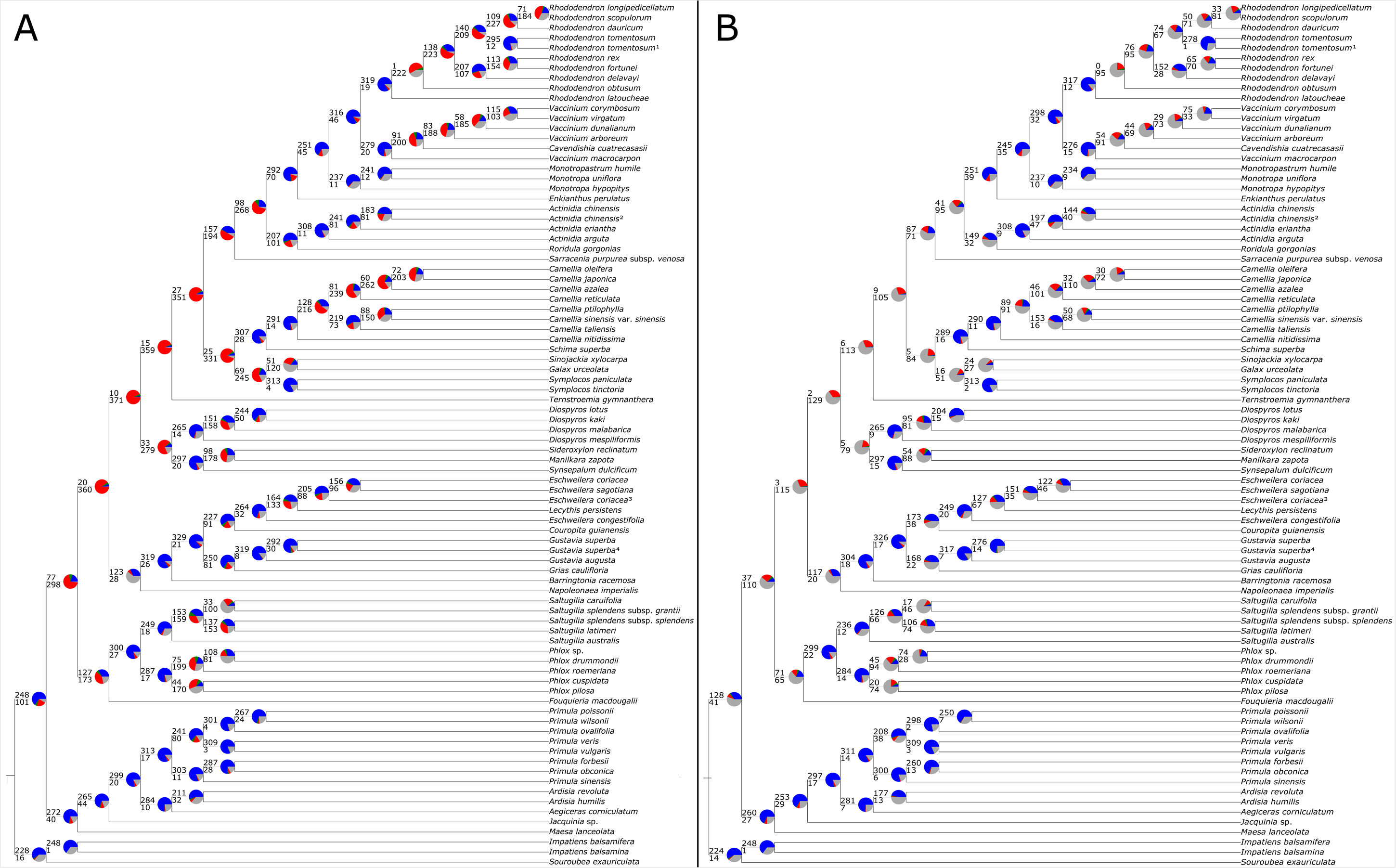
A) Ortholog tree concordance and conflict for the 387 ortholog set mapped to the rapid bootstrap topology. B) The same analysis except requiring 70% ortholog tree bootstrap support for an ortholog to be considered informative. Blue indicates the proportion of informative orthologs that are concordant with the topology, green indicates the proportion of informative orthologs that support the single most common conflicting topology, red indicates all other informative ortholog conflict and grey indicates orthologs that are uninformative, either due to support requirements or lack of appropriate taxon sampling.

**Supplemental Appendix 11.**
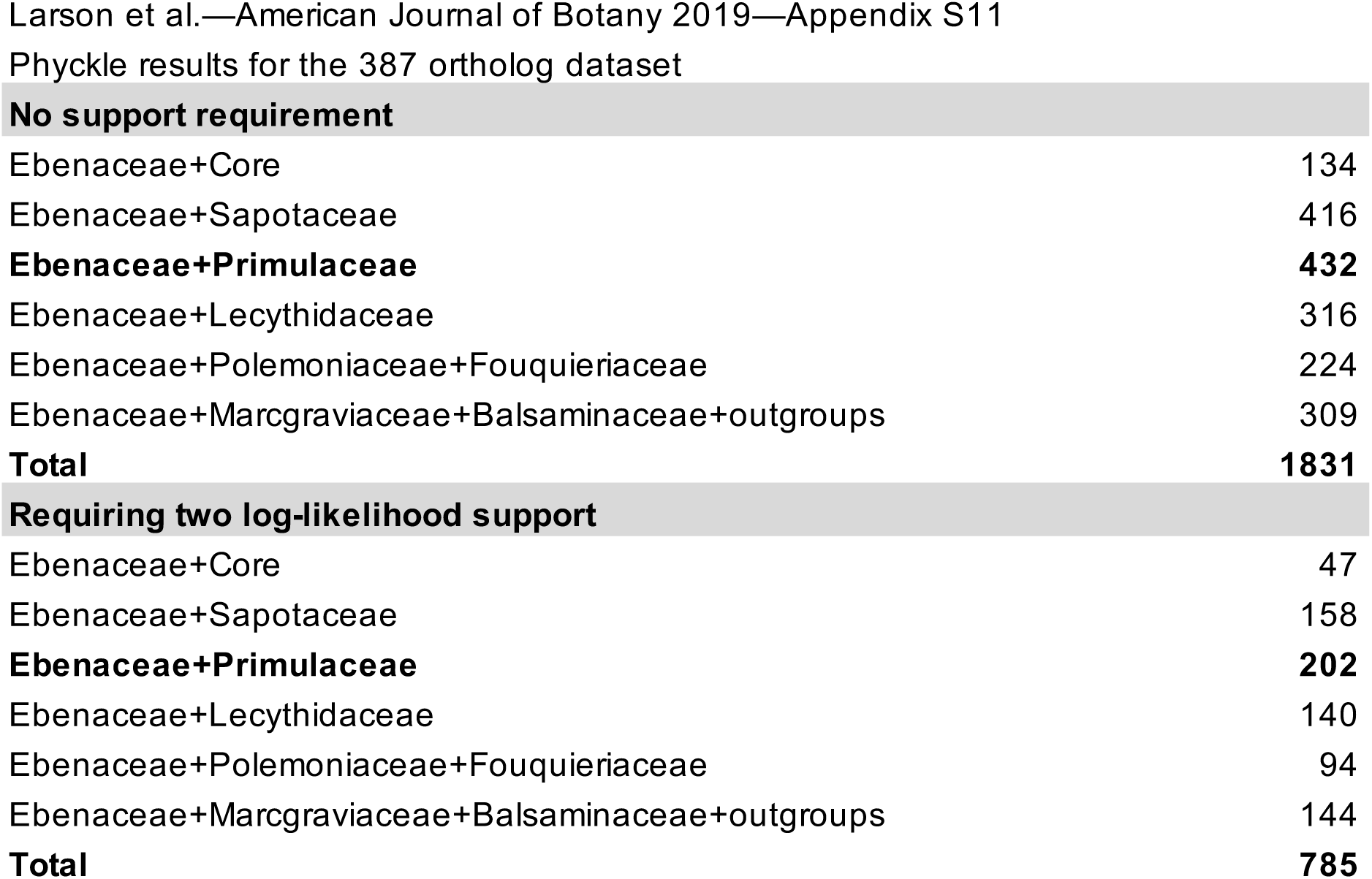
Phyckle results regarding the placement of Ebenaceae for the 2045 ortholog dataset. Ebenaceae+Primulaceae was supported by the highest number of orthologs whether or not two log-likelihood support difference was required.

**Supplemental Appendix 12.**
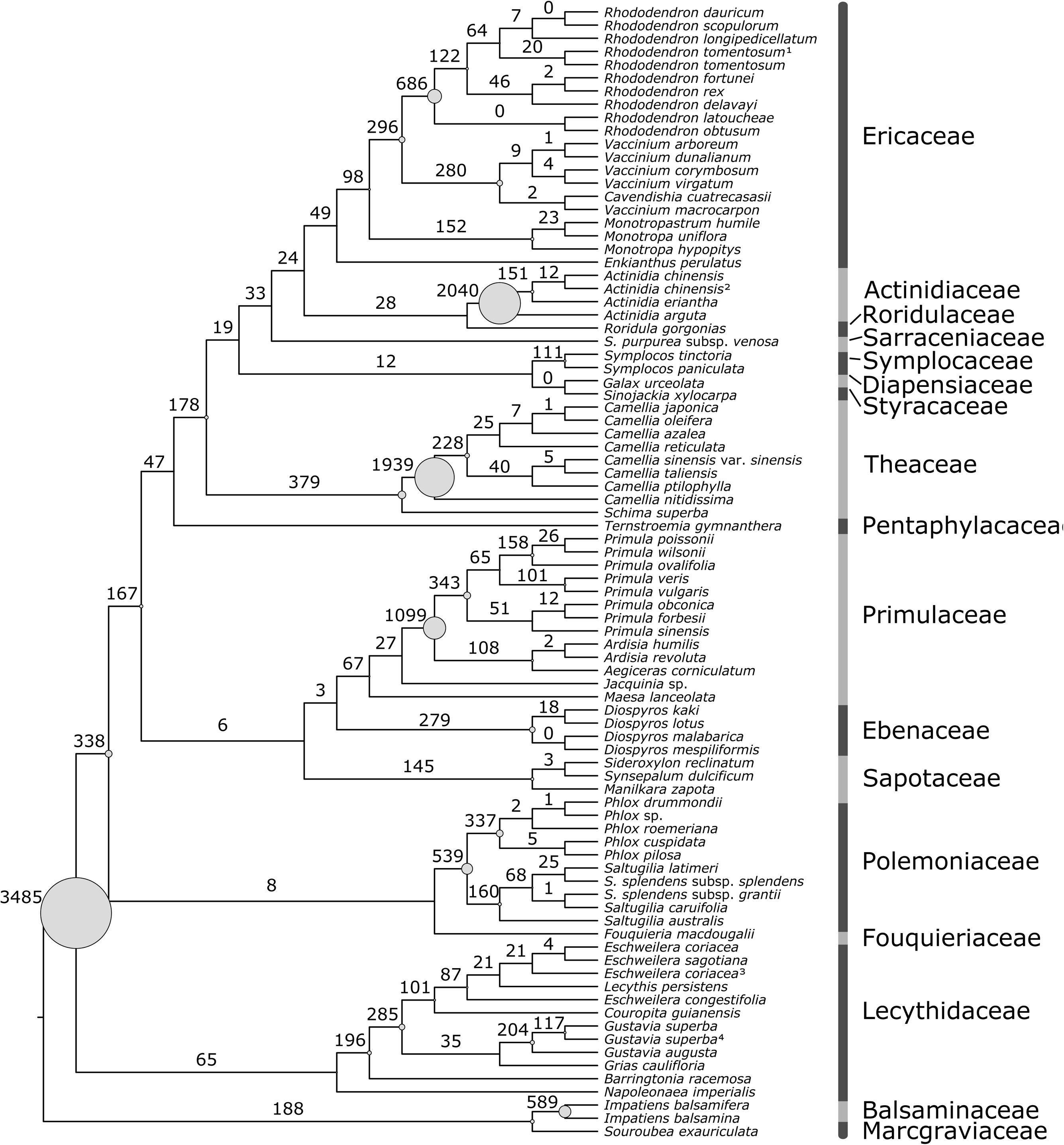
Gene duplications mapped to a cladogram of the 449 ortholog MAFFT-aligned, partitioned, supermatrix ML tree. Number of inferred gene duplications in shown along branches. The diameter of circles at nodes are proportional to the number of duplications.

**Supplemental Appendix 13.**
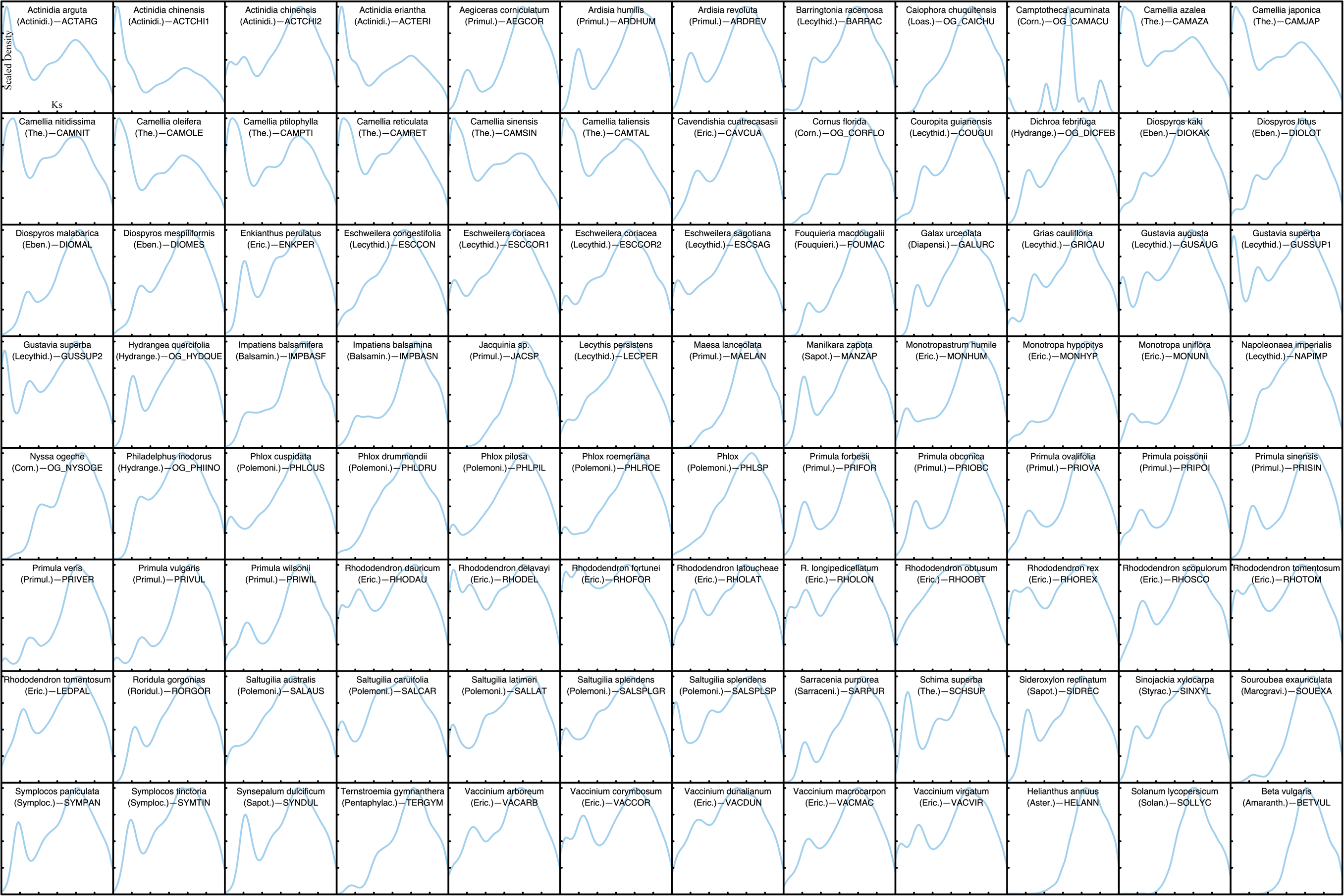
Single species K_s_ plots for pairs of paralogs within each taxa as a density plot. Density peaks indicate evidence for a large proportion of genes having been duplicated at approximately the same time, as would occur during a whole genome duplication. For each taxa, the x-axis ranges from 0.0 to 3.0 with each tick representing 0.5 synonymous substitutions between paralogs. The y-axis is scaled to the maximum density value for each transcriptome and ticks correspond to 0.25, 0.5, 0.75, and 1.0 of the maximum density respectively. Input transcriptomes have been filtered to remove short sequences and sample-specific duplications since these can represent transcript splice-site variants or errors in assembly.

**Supplemental Appendix 14.**
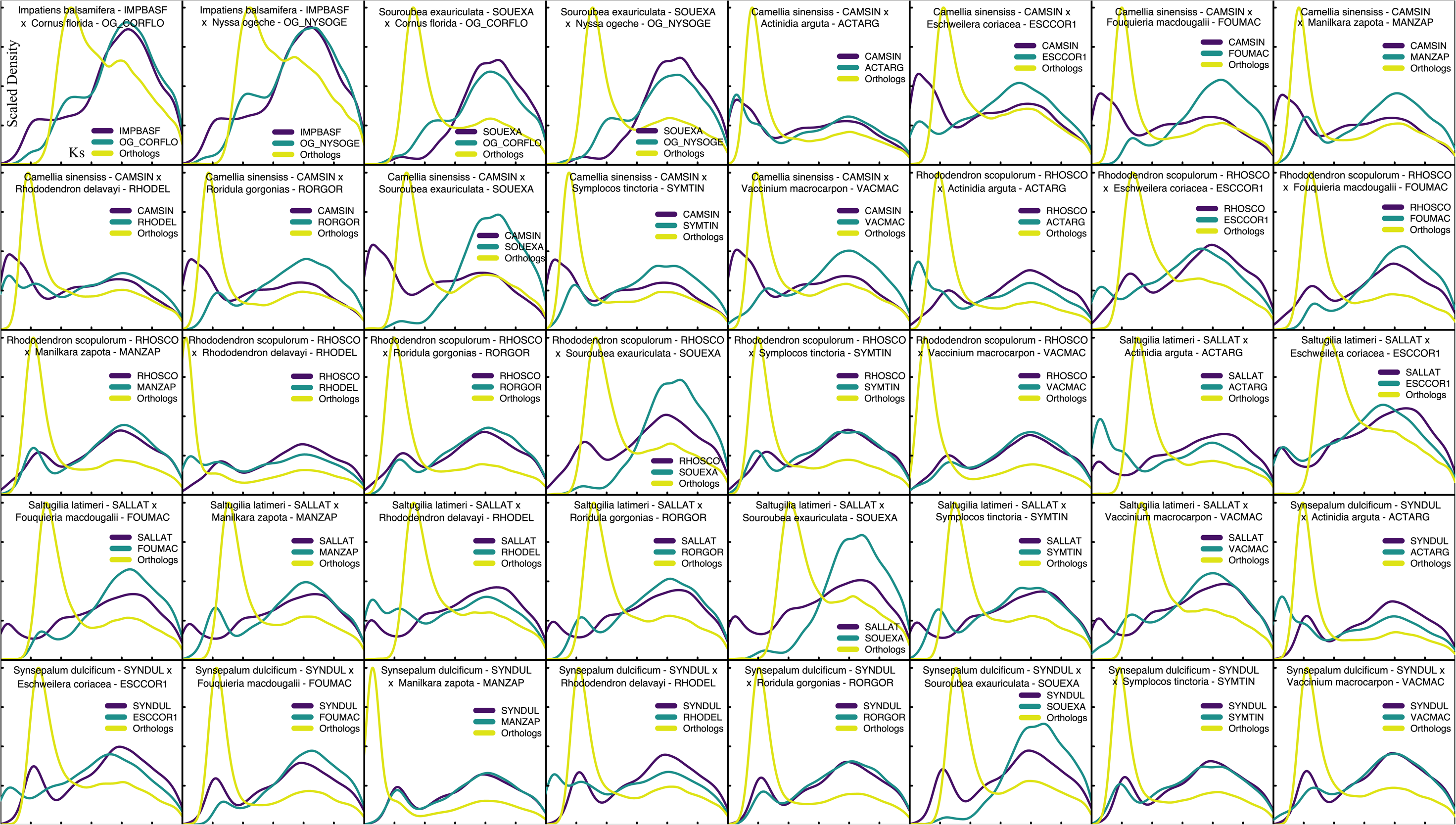
Multispecies K_s_ plots representing a broad range of taxonomic pairings. Single species K_s_ density plots (i.e. pairs of paralogs) within each taxa are plotted on the same axis as a K_s_ density plot representing pairs of orthologs between the two taxa. The density peak for the orthologs corresponds to the time of divergence between the two taxa, since orthologs in both taxa begin accumulating synonymous substitutions after speciation. If evolutionary rate in the two taxa are equal and the accumulation of synonymous substitutions is clock-like, then the timing the duplication events relative to divergence of the two taxa can be compared, with older events occurring farther to the right. The x-axis ranges from 0.0 to 3.0 with each tick representing 0.5 synonymous substitutions. The y-axis is scaled to the maximum density value of any of the three density distributions (therefore the magnitudes of all peaks are relative) and ticks correspond to 0.25, 0.5, 0.75, and 1.0 of the maximum density respectively. Input transcriptomes have been filtered to remove short sequences and sample-specific duplications since these can represent transcript splice-site variants or errors in assembly.

**Supplemental Appendix 15.**
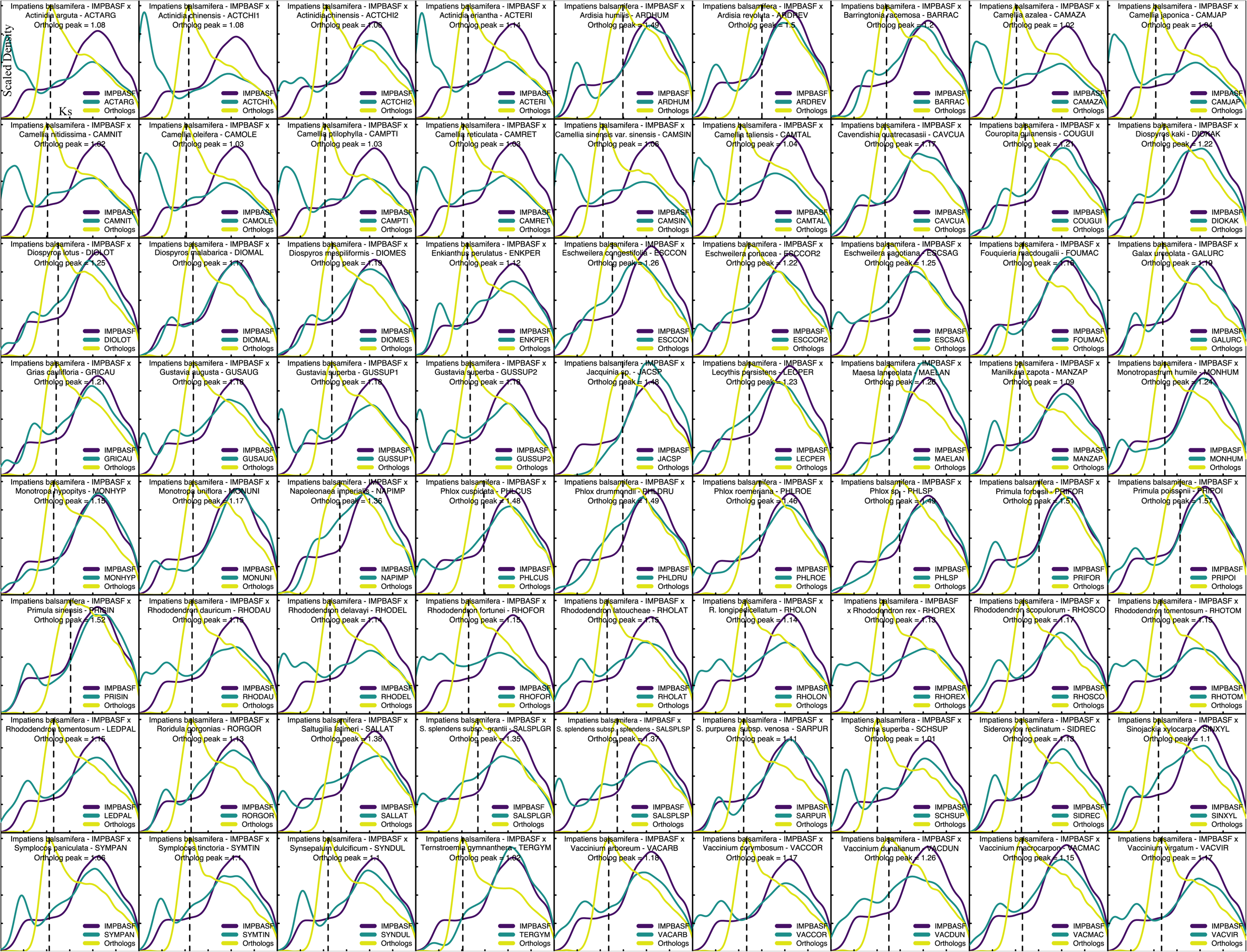
Multispecies K_s_ plots representing each non-balsaminoid ericalean taxa paired with *Impatiens balsamifera* for which there is an unambiguous ortholog peak. Single species K_s_ density plots (i.e. pairs of paralogs) within each taxa are plotted on the same axis as a K_s_ density plot representing pairs of orthologs between the two taxa. The density peak for the orthologs corresponds to the time of divergence between the two taxa, since orthologs in both taxa begin accumulating synonymous substitutions after speciation. The dashed line represents the point along the x-axis where the ortholog peak achieves its maximum value and would be expected to occur at approximately the same point in all pairings under clock-like accumulation of synonymous substitutions since all pairs share the same MRCA. The x-axis ranges from 0.0 to 3.0 with each tick representing 0.5 synonymous substitutions. The y-axis is scaled to the maximum density value of any of the three density distributions (therefore the magnitudes of all peaks are relative) and ticks correspond to 0.25, 0.5, 0.75, and 1.0 of the maximum density respectively. Input transcriptomes have been filtered to remove short sequences and sample-specific duplications since these can represent transcript splice-site variants or errors in assembly.

